# Machine Learning in Multi-Omics Data to Assess Longitudinal Predictors of Glycaemic Health

**DOI:** 10.1101/358390

**Authors:** Laurie Prélot, Harmen Draisma, Mila D. Anasanti, Zhanna Balkhiyarova, Matthias Wielscher, Loic Yengo, Beverley Balkau, Ronan Roussel, Sylvain Sebert, Mika Ala-Korpela, Philippe Froguel, Marjo-Riitta Jarvelin, Marika Kaakinen, Inga Prokopenko

**Affiliations:** Section of Genomics of Common Disease, Department of Medicine, Imperial College London, London W12 0NN, United Kingdom; Department of Epidemiology and Biostatistics, School of Public Health, Imperial College London, London W2 1NY, United Kingdom; Institute for Molecular Bioscience, The University of Queensland, 306 Carmody Road, St Lucia Qld 4072, Australia; Inserm, CESP Center for Research in Epidemiology and Public Health, U1018, Villejuif, France; Univ Paris-Saclay, Univ Paris Sud, UVSQ, UMRS 1018, UMRS 1018, Villejuif, France; Inserm U1138, Centre de Recherche des Cordeliers, Paris, France; University Paris Diderot, Sorbonne Paris Cite, Paris, France; Diabetology, Endocrinology and Nutrition Department, DHU FIRE, Hopital Bichat, AP-HP, Paris, France; Center for Life Course Health Research, University of Oulu, FI-90014 University of Oulu, Finland; Biocenter Oulu, University of Oulu, FI-90014 University of Oulu, Finland; Systems Epidemiology, Baker Heart and Diabetes Institute, Melbourne VIC 3004, Australia; Population Health Science, Bristol Medical School, University of Bristol, Bristol, UK; Medical Research Council Integrative Epidemiology Unit at the University of Bristol, Bristol BS8 1QU, United Kingdom; Computational Medicine, Faculty of Medicine, University of Oulu and Biocenter Oulu, FI-90014 University of Oulu, Finland; NMR Metabolomics Laboratory, School of Pharmacy, University of Eastern Finland, FI-70210 Kuopio, Finland; Department of Epidemiology and Preventive Medicine, School of Public Health and Preventive Medicine, Faculty of Medicine, Nursing and Health Sciences, The Alfred Hospital, Monash University, Melbourne VIC 3004, Australia; CNRS, Pasteur Institute of Lille, University of Lille, 59000 Lille, France; Oulu University Hospital, 90220 Oulu, Finland; Department of Life Sciences, College of Health and Life Sciences, Brunel University London, Uxbridge UB8 3PH, United Kingdom; Centre for Pharmacology and Therapeutics, Department of Medicine, Imperial College London, London W12 0NN, United Kingdom; School of Biosciences and Medicine, Department of Clinical and Experimental Medicine, University of Surrey, Guildford, United Kingdom

**Keywords:** Glycaemic traits, longitudinal, machine learning, metabolomics, methylation, prediction, type 2 diabetes

## Abstract

Type 2 diabetes (T2D) is a global health burden that will benefit from personalised risk prediction and targeted prevention programmes. Omics data have enabled more detailed risk prediction; however, most studies have focussed on directly on the ability of DNA variants predicting T2D onset with less attention given to epigenetic regulation and glycaemic trait variability. By applying machine learning to the longitudinal Northern Finland Birth Cohort 1966 (NFBC 1966) at 31 (T1) and 46 (T2) years old, we predicted fasting glucose (FG) and insulin (FI), glycated haemoglobin (HbA1c) and 2-hour glucose and insulin from oral glucose tolerance test (2hGlu, 2hIns) at T2 in 513 individuals from 1,001 variables at T1 and T2, including anthropometric, metabolic, metabolomic and epigenetic variables. We further tested whether the information obtained by the machine learning models in NFBC could be used to predict glycaemic traits in the independent French study with 48 matching predictors (DESIR, N=769, age range 30-65 years at recruitment, interval between data collections: 9 years). In this study, FG and FI were best predicted, with average R^2^ values of 0.38 and 0.53. Sex, branched-chain and aromatic amino acids, HDL-cholesterol, glycerol, ketone bodies, blood pressure at T2 and measurements of adiposity at T1, as well as multiple methylation marks at both time points were amongst the top predictors. In the validation analysis, we reached R^2^ values of 0.41/0.55 for FG/FI when trained and tested in NFBC1966 and 0.17/0.30 when trained in NFBC1966 and tested in DESIR. We identified clinically relevant sets of predictors from a large multi-omics dataset and highlighted the potential of methylation markers and longitudinal changes in prediction.

## Background

Diabetes accounts for the yearly deaths of about four million people between 20 and 79 years old (2017) world-wide and the prevalence of diabetes is expected to increase from 8.8% to 9.9% by 2045.^1^ Moreover, glucose tolerance impairment is progressing in young individuals, leading to high risk of developing type 2 diabetes (T2D) later in life.^1^

To date, T2D risk prediction in the clinical practice has focussed on the classical risk factors of sex, age, obesity, family history, hypertension, cholesterol levels and lifestyle factors ^2–5^. Recent advances in omics technologies have allowed exploring the risk factors of T2D in more detail, opening possibilities for more precise biomarkers and thus, better identification of people at risk for T2D in the future. A large number of metabolites, including amino acids, especially branched-chain amino acids (BRACA) and aromatic amino acids, fatty acids, glycerophospholipids, ketone bodies and mannose have been associated with T2D incidence^6–8^. However, whether metabolites can be effective and reliable T2D predictors, remains unclear.

During the past decade, genome-wide association studies (GWAS) have enlightened the genetic risk of developing T2D. Currently, 403 independent DNA variants are established for T2D risk by GWAS metaanalyses^9,10^ as well as dozens of loci have been associated with quantitative glycaemic traits in individuals without T2D, including fasting glucose (FG)^11^, fasting insulin (FI)^11^, FG adjusted for body-mass-index (BMI)^11^, FI adjusted for BMI^11^, 2 hour post-prandial or post oral glucose-tolerance test glucose (2hGluc) ^12^ and glycaeted haemoglobin (HbA1c)^13^. Besides genetics, environment and lifestyle are likely to have a large contribution to T2D risk^14^. Environmental factors can affect gene expression by an addition of a methyl group on a CpG-dinucleotide site of DNA. This is called DNA methylation and is the most widely studied type of epigenetic modification^15,16^. Studies in peripheral blood have found a mean absolute difference of 0.5-1.1% in methylation levels between individuals with and without T2D^17^. Epigenome-wide association studies have reported associations at 65 methylation markers for T2D^17,18^ and provided support for overlap in epigenetic effects between T2D and glycaemic traits^18,19^. The epigenetic effect on variability of glycaemic traits is magnified in the presence of obesity^19^. The investigation of the link between BMI and methylation levels demonstrates that methylation at the majority of CpG sites in blood is consequential to higher BMI^20^. Interestingly, a weighted methylation risk score calculated from 187 markers was shown to have an even stronger effect on incident T2D^20^ than the traditional risk factors of overweight, central obesity, phenylalanine, tyrosine, isoleucine, FG, FI and C-reactive protein. The methylation risk score remained associated with incident T2D even after adjustment for age, sex, BMI, FI, FG and central obesity^20^.

In the era of multi-omics data and millions of measurable variables, it is challenging to identify the best biomarkers for clinical use. Machine learning approaches are ideal for such high-dimensional data due to their ability to learn from the data and identify patterns without knowledge of the joint distribution of the variables^21^. Recently, the studies of T2D risk have leveraged association analyses as well as machine learning algorithms for the prediction of binary T2D phenotypes. Thus far, machine learning used for T2D classification include logistic regression with and without Lasso regularization^17,20,22–26^, Regularized least-squares (RLS)^27^, Cox regression^23^, naïve Bayes^25^ and J48-decision tree^25^. In predictive studies using machine learning models, classical risk factors, genetic risk scores (GRS)^22^, methylation risk scores (MRS)^17,20^ and metabolomic data^24–27^ have been used as predictors of T2D incidence after a follow-up window of two to fourteen years. A few studies have suggested that metabolites improve prediction performance^23,25–27^, while others have reported negligible to no improvement in prediction^24^. GRS have been shown to bring no incremental value over classical non-invasive factors and metabolic markers^22^. MRS combining CpG loci have been found to be associated with future T2D incidence^17,20^. All the previous studies have focused on the binary disease status as the outcome which may greatly reduce the power of the analyses. To our knowledge, there are no machine learning studies on predictors of continuous glycaemic traits relevant for T2D pathophysiology. In addition, most of the studies on T2D have used a single pre-selected machine learning approach and have not compared their performance in terms of predictive capacities. With the present study, we aimed to address these shortcomings, as well as to shed light on the contribution of longitudinal predictors, especially metabolomic and methylation markers by using data from two well-characterised population-based studies and by applying and comparing six different machine learning approaches.

## Results

We focused on epigenetic and metabolic markers (**Supplementary Table 1, Supplementary Table 2**) from the Northern Finland Birth Cohort 1966 (NFBC1966) measured at 31 (T1) and 46 (T2) years for prediction of HbA1c, FG, 2hGluc, FI and two-hours post-oral glucose tolerance test insulin (2hIns) at T2 in individuals free from T2D diagnosis or medication at T1. We implemented and compared the results obtained from six machine learning approaches: Boosted trees (BT), Random forest (RF) and support vector regression (SVR) with Linear Kernel with L2 regularization and with L1 and L1/L2 loss functions (SVR-L2Linear-L1, SVR-L2Linear-L1L2, respectively), with Polynomial Kernel (SVR-Polynomial) and with Radial Basis function Kernel (SVR-RBF). The algorithms were chosen for their ability to handle a large number of predictors, to account for multi-collinearity, non-linear relationships, the absence of the assumption regarding data distribution, and for their computational times. We also tested different input data combinations (Figure 1). We further validated our approach in an independent French cohort (Data from an Epidemiological Study on the Insulin Resistance syndrome, DESIR) that shared 48 variables with the NFBC1966 cohort (**Supplementary Table 3**).

**Figure 1.**
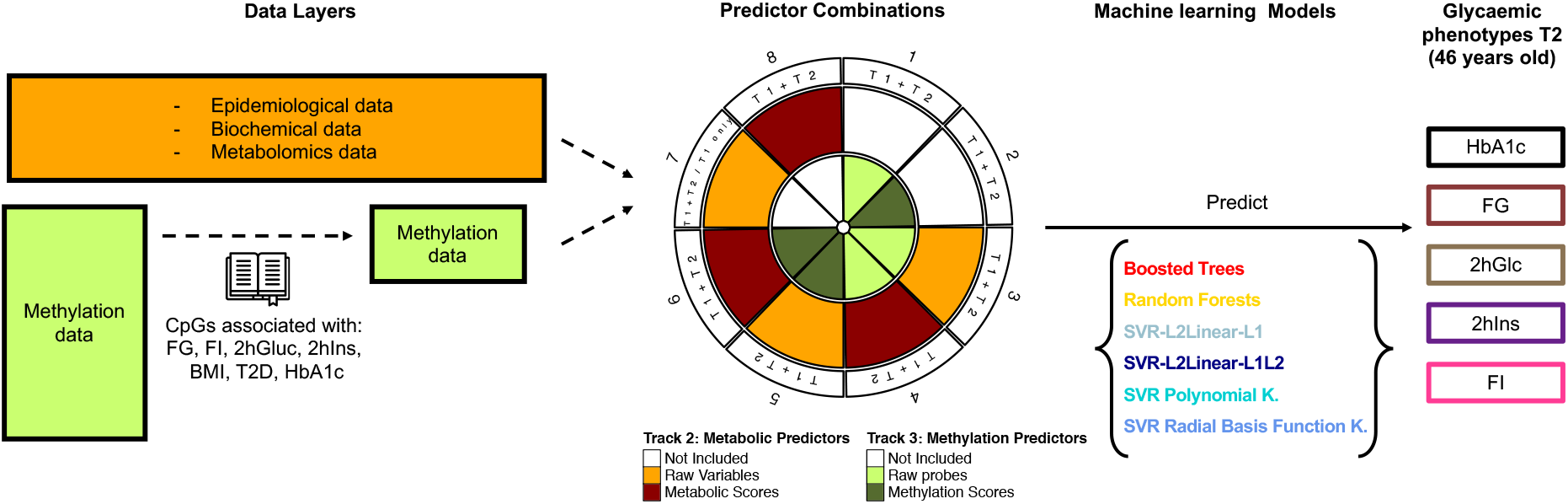
Experimental set-up for Machine learning analysis. We applied machine learning to multiomics data based on blood samples and data collections from the Northern Finland Birth Cohort 1966 at 31 and 46 years. Fasting glucose/insulin (FG/FI), glycated haemoglobin (HbA1c) and 2-hour glucose/insulin (2hGlu/2hIns) phenotypes at T2 were predicted in 513 individuals using up to 991 variables from T1 and T2: Body-mass-index (BMI), waist-hip-ratio, systolic and diastolic blood pressure (SBP and DBP), sex; 10 blood plasma measurements; 453 NMR-based metabolites; 528 methylation probes established for BMI, FG, FI, HbA1c, 2hGlu, 2hIns or Type 2 diabetes. Six machine learning approaches were used: random forest, boosted trees and support vector regression (SVR) with the kernels of linear function with L2 regularization and L1 loss function (L2Linear-L1), linear with L2 regularization and L1/L2 loss function (L2Linear-L1L2), polynomial and radial basis function (RBF).

### Best predicted outcome variables

We compared the performance of the six models for each of the outcomes (HbA1c, FG, 2hGluc, FI and 2hIns) with varying input data combinations, including omics data in their raw or scored forms (**Methods, Supplementary Table 4**). With the usage of metabolic data as the minimal input, we observed that performance was the best for FI prediction (Figure 2A). In the context of the model with metabolic raw data (Mb-R), models with different outcomes were ranked as follows: FI > FG > 2hIns > 2hGlc > HbA1c (*P*_TukeyHSD_<5.0×10^-5^ for all other comparisons except for 2hIns>2hGluc for which *P*_TukeyHSD_=0.19, **Supplementary Table 5**). The average coefficients of determination R^2^ over all machine learning algorithms were 0.53, 0.38, 0.29, 0.25 and 0.14 (Mb-R) for FI, FG, 2hIns, 2hGlc, HBA1c respectively (**Supplementary Table 6**).

**Figure 2.**
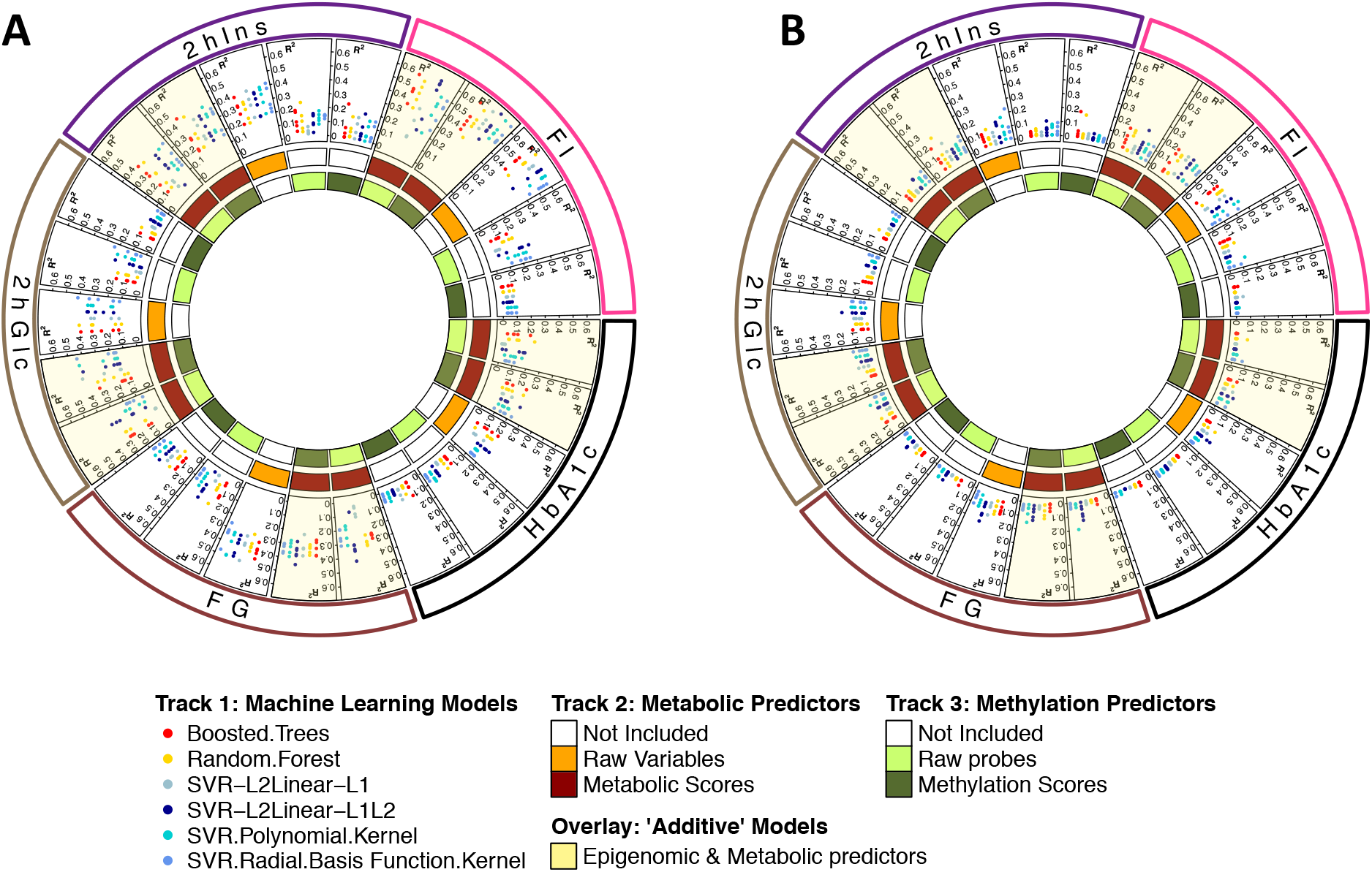
Performance as measured in R^2^ of the different machine learning models. A. Unadjusted for measurements of adiposity (Waist-to-hip-ratio and Body mass index) at T1 and T2; B. Adjusted for measurements of adiposity (Waist-to-hip-ratio and Body mass index) at T1 and T2. Training of the algorithm was performed with a nested cross validation (5-folds outer, and 5-folds inner cross validation) and the R^2^ of 5 outer testing folds is displayed for each machine learning model. Metabolic predictors include epidemiological, biochemical and metabolomic data. SVR: Support Vector Regression with the kernels of linear function with L2 regularization and L1 loss function (L2Linear-L1), linear with L2 regularization and L1/L2 loss function (L2Linear-L1L2), polynomial and radial basis function (RBF).

### Best predictor data combinations

Regarding the predictors, we found that models that included at least some metabolic data, either in their raw format (Mb-R) or transformed into scores (Mb-S) (**Methods**) had the best performance reaching a maximum R^2^=0.56 (Figure 2A, **Supplementary Table 7**). In contrast, models with methylation data only as predictors (Mh-R and Mh-S), reached R^2^ values of up to 0.20. Thus, metabolic models performed significantly better than pure epigenomic models (*P*_TukeyHSD_<1.7×10^-9^) (Table 1, Comparison 1-4; all comparisons are provided in **Supplementary Table 8**). When metabolic and methylation data were combined, Mb-R performed better for FI and FG than Mb-R + Mh-R (*P*_TukeyHSD_ <0.04). Adding Mh-S to the model did not alter the model performance for any of the outcomes (*P*_TurkeyHSD_>0.99) (Table 1, Comparison 5-6). These results suggest that addition of methylation information does not increase the predictive ability of the tested models. When exploring the effect of transforming the original variables into scores (Table 1, comparison 7-8), we observed no significant differences for any of the outcomes when comparing Mb-R vs. Mb-S (*P*_TukeyHSD_>0.29). This was true also for Mh-R vs. Mh-S, except for the prediction of FI, where Mh-R performed better than Mh-S (*P*_TukeyHSD_=1.6×10^-4^). Therefore, based on these results we are unable to generalise better performance of the scored data as compared to raw data. Finally, we found that Mb-S + Mh-S model performed significantly better than the model with Mb-S + Mh-R (*P*_TukeyHSD_<0.02) for all outcomes except for 2hGluc. This observation reflects the decrease in performance of the models upon inclusion of a large number of weak predictors.

**Tables 1.**
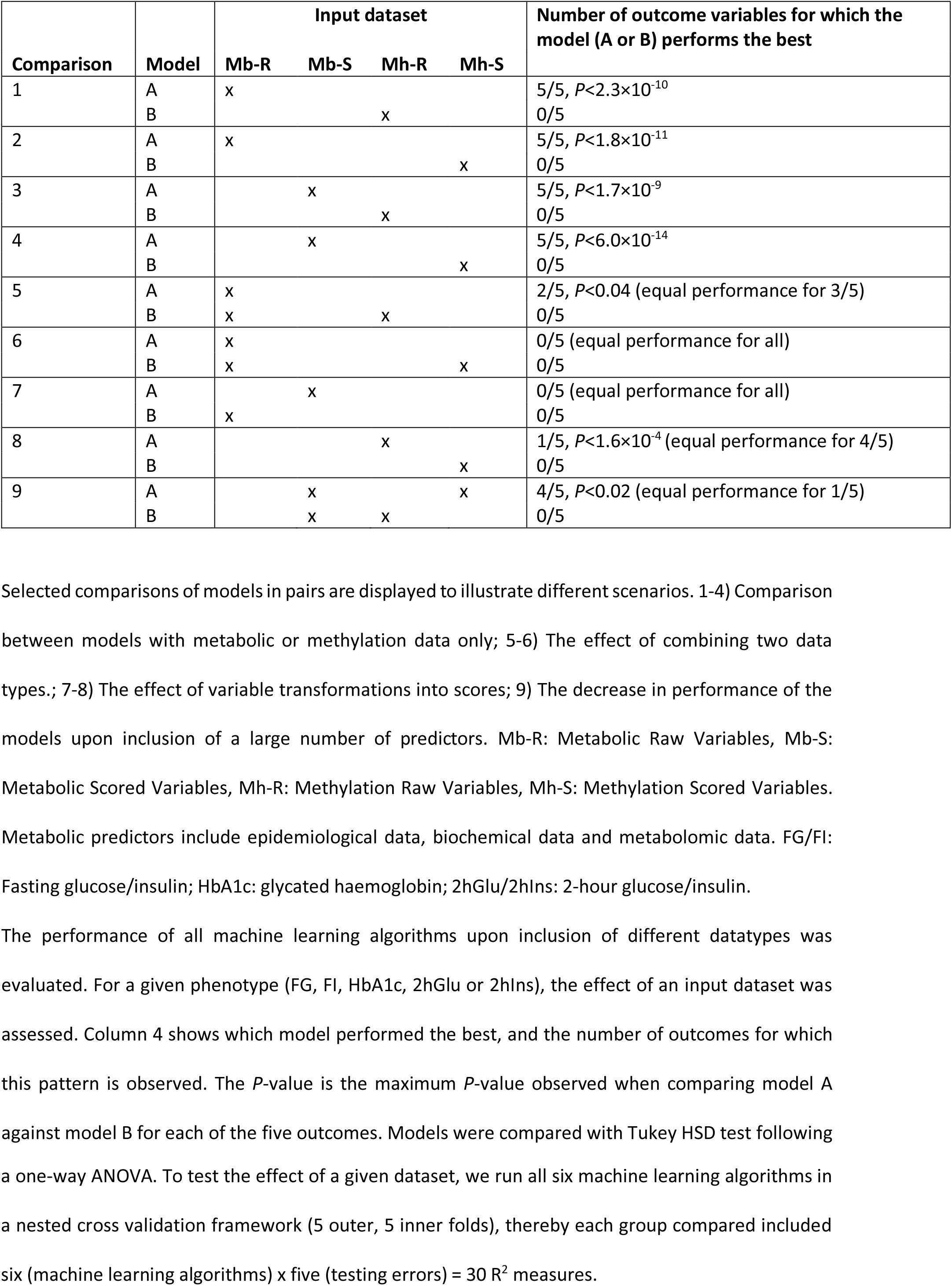
Effect of the input dataset on the prediction performance of the five glycaemic traits.

### Adjustment for Measures of Adiposity

To understand the influence of the measures of adiposity in the models, we adjusted all the outcomes for T1-BMI, T2-BMI, T1-WHR and T2-WHR and performed the machine learning models on these adjusted outcomes. All adjusted models exhibited an R^2^<0.13 (Figure 2B, **Supplementary Figure 1** with a zoomed-in scale for R^2^, **Supplementary Table 9**), including models predicting FI and FG. Therefore, the measures of adiposity at T1 and T2 are the main drivers of prediction for FI and FG.

### Variable Importance

We investigated the contribution of metabolic and epigenomic variables to the prediction of glycaemic traits. We discuss predictors importance only in the context of FG and FI outcomes, for which prediction algorithms reached the best R^2^ (Figure 2A). FG prediction was mostly explained by metabolic variables: valine, leucine, isoleucine, tyrosine, BMI and WHR, HDLs and VLDL, glycerol, alanine, SBP and DBP at T2, WHR and FG at T1 and sex (Figure 3 bottom right). FI prediction was explained by BMI, WHR, HDL, VLDL, BRACA, phenylalanine, leucine, glycerol, lactate, tyrosine, valine at T2, and FI at T1 (Figure 3 top left). Once we adjusted the outcomes for the measures of adiposity, the top predictor for FI was measurement of FI at T1, followed by alanine and other variables at T2 already observed as important before adjustment for measurements of adiposity (**Supplementary Figure 2**). The top variables driving the prediction of FG were, similarly to FI, valine at T2 and FG at T1 (**Supplementary Figure 2**). The metabolic models with scored variables were driven by variables that mirrored the top raw predictors. Overall, the model with scored variables for FI supported the importance of the former variables, as well as ketone bodies (acetoacetate and 3-hydroxybutyrate) at T2 (Figure 3, top right and bottom left). Importantly, even though we previously observed that adding methylation data does not improve model prediction, a number of methylation probes were among the top predictors for both FG and FI when combined with scored metabolic variables (Figure 3).

**Figure 3.**
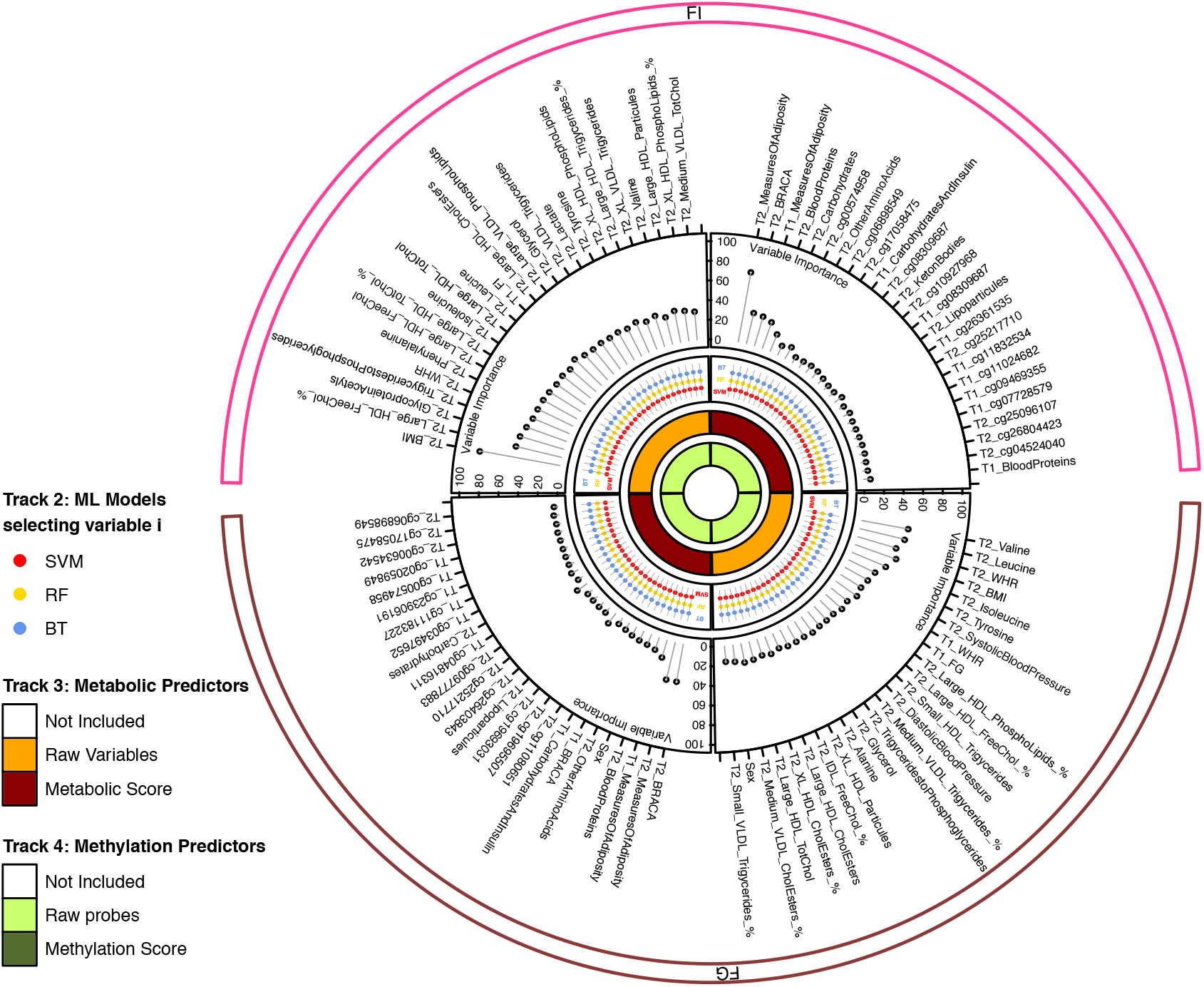
Variable Importance for fasting glucose and fasting insulin prediction from two different combinations of predictor data. For each Machine learning method, the normalized variable importance over five outer fold of cross validation was averaged into the “Variable-Model-Importance” (var.mod.Imp). Then for each of the six machine learning models, the variables were ranked based on the var.mod.Imp. The rank was averaged over the six models to obtain the “mean variable rank”. The latter was used to select top 25 variables for display. For these variables, we display the variable importance after (1) weighting the var.mod.Imp by the R^2^ obtained for each of the individual machine learning algorithms (2) averaging variable importance across the six machine learning models. FI: Fasting Insulin; FG: Fasting Glucose; RF: random forest; BT: boosted trees, SVR: support vector regression models with the kernels of linear function with L2 regularization and L1 loss function (L2Linear-L1), linear with L2 regularization and L1/L2 loss function (L2Linear-L1L2), polynomial and radial basis function (RBF). Metabolic predictors include epidemiological data, biochemical data and metabolomic data. T2: 46 years old, T1: 31 years old. BMI: Body Mass Index according to clinical examination, postal questionnaire if missing; WHR: Waist-to-hip ratio. Metabolite name descriptions are provided in **Supplementary Table 1**.

### Association Analysis for Effect Sizes, Direction and Variance Explained

Linear regression analyses between ln(FI)/FG and each of the top 25 predictors identified by machine learning indicated that increases in HDL cholesterol were predictive of decreased values of FI and FG (Table 2). The same was true for free cholesterol to total lipids ratio in IDL and cholesterol esters to total lipids ratio in medium VLDL. Similarly, female gender and higher sum score of ketone bodies, i.e. acetoacetate and 3-hydroxybutyrate, predicted lower FI and FG levels, as did higher methylation levels of the probes T2_cg00574958, T2_cg17058475, and T2_cg08309687. In the models with scored variables, the adjusted R^2^ for ln(FI) when including all 25 top predictors was 0.58. After exclusion of methylation markers, it decreased to 0.54 and finally, after excluding all other variables than measures of adiposity, the variance explained was 0.40. The same figures for FG were 0.45, 0.42 and 0.31, respectively.

**Table 2.** Linear regression analysis results for In(FI)/FG and the top 25 predictors.

| **Mb-R + Mh-R** | **beta** | **SE** | **P-value** | **Mb-S + Mh-R** | **beta** | **SE** | **P-value** |
| --- | --- | --- | --- | --- | --- | --- | --- |
| **ln(FI)** |  |  |  | **ln(FI)** |  |  |  |
| T2_BMI | 0.08 | 0.00 | $3.29 \times 10^{-55}$ | T2_MeasuresOfAdiposity | 0.08 | 0.00 | $5.75 \times 10^{-56}$ |
| T2_Large_HDL_FreeChol_% | -0.11 | 0.01 | $1.48 \times 10^{-40}$ | T2_BRACA | 3.76 | 0.29 | $1.17 \times 10^{-32}$ |
| T2_GlycoproteinAcetyls | 1.32 | 0.10 | $5.88 \times 10^{-37}$ | T1_MeasuresOfAdiposity | 0.07 | 0.01 | $7.39 \times 10^{-26}$ |
| T2_TriglyceridestoPhosphoglycerides | 1.13 | 0.09 | $8.73 \times 10^{-35}$ | T2_BloodProteins | 1.32 | 0.10 | $3.36 \times 10^{-37}$ |
| T2_WHR | 3.79 | 0.27 | $3.12 \times 10^{-38}$ | T2_Carbohydrates | 0.73 | 0.06 | $1.62 \times 10^{-26}$ |
| T2_Phenylalanine | 29.08 | 2.32 | $1.62 \times 10^{-31}$ | T2_cg00574958 | -9.28 | 1.48 | $8.62 \times 10^{-10}$ |
| T2_Large_HDL_FreeChol | -4.91 | 0.39 | $1.03 \times 10^{-31}$ | T2_OtherAminoAcids | 0.66 | 0.18 | $3.42 \times 10^{-4}$ |
| T2_Large_HDL_TotChol_% | -0.04 | 0.00 | $2.04 \times 10^{-34}$ | T2_cg06898549 | 3.01 | 0.63 | $2.54 \times 10^{-6}$ |
| T2_Isoleucine | 15.32 | 1.23 | $2.52 \times 10^{-31}$ | T2_cg17058475 | -4.67 | 1.12 | $3.45 \times 10^{-5}$ |
| T2_Large_HDL_TotChol | -1.14 | 0.09 | $4.61 \times 10^{-30}$ | T1_CarbohydratesAndInsulin | 0.10 | 0.02 | $1.68 \times 10^{-9}$ |
| T2_Leucine | 13.13 | 1.07 | $1.09 \times 10^{-30}$ | T2_cg08309687 | -3.51 | 0.68 | $3.24 \times 10^{-7}$ |
| T1_FI | 0.80 | 0.07 | $1.19 \times 10^{-27}$ | T2_KetonBodies | -0.70 | 0.29 | $1.52 \times 10^{-2}$ |
| T2_Large_HDL_CholesterolEsters | -1.47 | 0.12 | $1.86 \times 10^{-29}$ | T2_cg10927968 | 3.01 | 0.64 | $3.23 \times 10^{-6}$ |
| T2_Large_VLDL_PhosphoLipids | 4.59 | 0.41 | $1.76 \times 10^{-26}$ | T1_cg08309687 | -2.44 | 0.63 | $1.10 \times 10^{-4}$ |
| T2_Glycerol | 11.03 | 0.94 | $3.94 \times 10^{-28}$ | T2_Lipoparticules | 0.00 | 0.00 | $4.99 \times 10^{-9}$ |
| T2_VLDL_Triglycerides | 0.40 | 0.03 | $6.03 \times 10^{-28}$ | T1_cg26361535 | 1.76 | 0.70 | $1.29 \times 10^{-2}$ |
| T2_Lactate | 0.74 | 0.07 | $2.34 \times 10^{-25}$ | T2_cg25217710 | 4.83 | 1.27 | $1.51 \times 10^{-4}$ |
| T2_Tyrosine | 20.79 | 1.87 | $5.98 \times 10^{-26}$ | T1_cg11832534 | 3.30 | 1.13 | $3.54 \times 10^{-3}$ |
| T2_XL_HDL_PhosphoLipids | -1.30 | 0.11 | $1.81 \times 10^{-28}$ | T1_cg11024682 | 3.92 | 1.02 | $1.33 \times 10^{-4}$ |
| T2_Large_HDL_Triglycerides_% | 0.10 | 0.01 | $3.53 \times 10^{-26}$ | T1_cg09469355 | -3.98 | 0.93 | $2.11 \times 10^{-5}$ |
| T2_XL_VLDL_Triglycerides | 4.04 | 0.38 | $3.47 \times 10^{-24}$ | T1_cg07728579 | 1.87 | 1.00 | $6.18 \times 10^{-2}$ |
| T2_Valine | 5.83 | 0.54 | $1.48 \times 10^{-24}$ | T2_cg25096107 | -3.36 | 1.40 | $1.68 \times 10^{-2}$ |
| T2_Large_HDL_Particules | -375767.70 | 31515.32 | $4.35 \times 10^{-29}$ | T2_cg26804423 | 4.02 | 1.10 | $2.96 \times 10^{-4}$ |
| T2_XL_HDL_PhosphoLipids_% | -0.02 | 0.00 | $2.03 \times 10^{-26}$ | T2_cg04524040 | -3.21 | 0.89 | $3.59 \times 10^{-4}$ |
| T2_Medium_VLDL_TotChol | 2.45 | 0.22 | $6.10 \times 10^{-26}$ | T1_BloodProteins | 0.39 | 0.08 | $1.52 \times 10^{-6}$ |
| **FG** |  |  |  | **FG** |  |  |  |
| T2_Valine | 7.43 | 0.56 | $7.64 \times 10^{-35}$ | T2_BRACA | 4.52 | 0.31 | $2.79 \times 10^{-41}$ |
| T2_Leucine | 15.38 | 1.12 | $1.26 \times 10^{-36}$ | T2_MeasuresOfAdiposity | 0.06 | 0.01 | $2.57 \times 10^{-28}$ |
| T2_WHR | 3.73 | 0.30 | $4.59 \times 10^{-31}$ | T1_MeasuresOfAdiposity | 0.06 | 0.01 | $2.56 \times 10^{-14}$ |
| T2_BMI | 0.06 | 0.01 | $8.60 \times 10^{-28}$ | T2_BloodProteins | 1.05 | 0.11 | $1.42 \times 10^{-19}$ |
| T2_Isoleucine | 16.56 | 1.33 | $2.53 \times 10^{-31}$ | Sex | -0.41 | 0.05 | $1.37 \times 10^{-14}$ |
| T2_Tyrosine | 20.74 | 2.05 | $5.73 \times 10^{-22}$ | T2_OtherAminoAcids | 0.96 | 0.19 | $1.08 \times 10^{-6}$ |
| T2_SystolicBloodPressure | 0.01 | 0.00 | $2.82 \times 10^{-20}$ | T1_BRACA | 1.88 | 0.31 | $4.10 \times 10^{-9}$ |
| T1_WHR | 2.80 | 0.30 | $7.01 \times 10^{-19}$ | T1_CarbohydratesAndInsulin | 0.13 | 0.02 | $4.46 \times 10^{-13}$ |
| T1_FG | 0.48 | 0.05 | $1.85 \times 10^{-18}$ | T2_cg11080651 | -8.53 | 1.94 | $1.39 \times 10^{-5}$ |
| T2_Large_HDL_PhosphoLipids_% | 0.05 | 0.01 | $1.04 \times 10^{-19}$ | T2_cg19695507 | 4.06 | 1.13 | $3.57 \times 10^{-4}$ |
| T2_Large_HDL_FreeChol_% | -0.09 | 0.01 | $2.80 \times 10^{-20}$ | T2_cg19693031 | -2.79 | 0.73 | $1.53 \times 10^{-4}$ |
| T2_Small_HDL_Triglycerides | 10.63 | 1.52 | $7.58 \times 10^{-12}$ | T2_Lipoparticules | 0.00 | 0.00 | $9.06 \times 10^{-8}$ |
| T2_DiastolicBloodPressure | 0.02 | 0.00 | $1.00 \times 10^{-16}$ | T2_cg26403843 | 2.07 | 0.60 | $6.44 \times 10^{-4}$ |
| T2_Medium_VLDL_Triglycerides_% | 0.03 | 0.00 | $7.61 \times 10^{-12}$ | T2_cg25217710 | 4.17 | 1.38 | $2.58 \times 10^{-3}$ |
| T2_TriglyceridestoPhosphoglycerides | 0.88 | 0.10 | $1.77 \times 10^{-17}$ | T2_cg09777883 | 2.20 | 1.09 | $4.32 \times 10^{-2}$ |
| T2_Glycerol | 8.74 | 1.08 | $4.71 \times 10^{-15}$ | T1_cg04816311 | 1.75 | 0.67 | $8.93 \times 10^{-3}$ |
| T2_Alanine | 2.89 | 0.35 | $8.41 \times 10^{-16}$ | T2_Carbohydrates | 0.51 | 0.07 | $2.37 \times 10^{-11}$ |
| T2_XL_HDL_Particules | -768070.10 | 98342.58 | $3.27 \times 10^{-14}$ | T1_cg03497652 | 1.96 | 0.70 | $5.47 \times 10^{-3}$ |
| T2_IDL_FreeChol_% | -0.18 | 0.02 | $2.22 \times 10^{-15}$ | T1_cg11183227 | 3.85 | 1.10 | $5.01 \times 10^{-4}$ |
| T2_Large_HDL_CholesterolEsters | -1.15 | 0.14 | $3.18 \times 10^{-15}$ | T1_cg23906191 | 4.99 | 1.89 | $8.39 \times 10^{-3}$ |
| T2_XL_HDL_CholesterolEsters_% | 0.02 | 0.00 | $7.27 \times 10^{-12}$ | T1_cg00574958 | -4.21 | 1.64 | $1.06 \times 10^{-2}$ |
| T2_Large_HDL_TotChol | -0.89 | 0.11 | $9.31 \times 10^{-16}$ | T1_cg02059849 | 5.61 | 1.36 | $4.20 \times 10^{-5}$ |
| T2_Medium_VLDL_CholesterolEsters_% | -0.04 | 0.01 | $8.94 \times 10^{-13}$ | T2_cg00634542 | 2.90 | 1.30 | $2.65 \times 10^{-2}$ |
| Sex | -0.41 | 0.05 | $1.37 \times 10^{-14}$ | T2_cg17058475 | -3.86 | 1.22 | $1.60 \times 10^{-3}$ |
| T2_Small_VLDL_Triglycerides_% | 0.03 | 0.00 | $1.58 \times 10^{-12}$ | T2_cg06898549 | 2.47 | 0.69 | $3.89 \times 10^{-4}$ |

The models were fitted for each predictor separately. Left panel shows the results when the predictors were in their raw format (Metabolic-Raw, Mb-R + Methylation-Raw, Mh-R) and right panel shows when the metabolic predictors were transformed into scores but methylation data kept in raw format (Metabolic-Score, Mb-S + Methylation-Raw, Mh-R).

### Prediction from Variables at T1

To test whether variables already 15 years beforehand, i.e. at T1 only, can provide information about glycaemic traits at T2, we restricted the prediction variables to those measured at T1 only. For example, under the well-performing Mb-R in the full model with predictors both from T1 and T2 and unadjusted for measures of adiposity, FI, FG, 2hIns, 2hGlc, HBA1c were predicted with R^2^ values of 0.53, 0.38, 0.29, 0.25 and 0.14, respectively. The restriction to T1 variables caused a drop in R^2^ to 0.25, 0. 22, 0.15, 0.06, 0.06, respectively, when averaging over all machine learning methods. This suggests that prediction from T1 variables only is not achievable in our dataset.

### Performance of the machine learning algorithms

When at least metabolic data in either form (Mb-R or –S + any other data) were included as input, we found no statistically significant differences between the performances of RF and BT for all phenotypes (*P*_TukeyHSD_>0.91, **Supplementary Table 10**). In addition, no significant difference was found between SVR-L2Linear-L1L2 and RF or BT models (*P*_TukeyHSD_>0.29, **Supplementary Table 10**). Among SVR models, we found that SVR-L2Linear-L1L2 either performed equally or outperformed the other SVRs, depending on the input dataset. In particular, for datasets with a large number of predictors SVR-L2Linear-L1L2 was the best performing SVR (*P*_TukeyHSD_<0.05, **Supplementary Table 10**). SVR-L2Linear-L1 in turn showed lower performance (*P*_TukeyHSD_<0.05, **Supplementary Table 10**) at several occasions when compared to that of the other algorithms. Even though both SVR-L2Linear-L1 and SVR-L2Linear-L1L2 both use the L2 regularization, the first uses only the L1 loss function whereas the latter optimizes over both L1 and L2 loss functions (**Supplementary Material**). We observed that in 78.5% of the time the L2 loss function was chosen over L1 in the SVR-L2Linear-L1L2 analyses (data not shown). We investigated whether the better performance of the SVR-L2Linear-L1L2 algorithm over the SVR-L2Linear-L1 was due to the evaluation criterion used, namely R^2^ which computes a scaled measure based on the quadratic loss function. For this purpose, we assessed the model performance in terms of Mean Absolute Error (MAE), i.e. based on L1 loss. By definition we aim to maximize R^2^ while we aim to minimize MAE. Evaluations based on MAE did not show improved performance of the predictions based on SVR-L2Linear-L1 (**Supplementary Figure 3, Supplementary Table 11**).

### Validation of the Machine Learning models in the French DESIR cohort

For the replication analysis, we first trained and tested the data in the NFBC1966 using the set of 48 variables common to both NFBC1966 and DESIR. Next, we trained the data in the NFBC1966 and tested in DESIR. We predicted only FI and FG levels using the top three performing algorithms: RF, BT and SVR-L2Linear-L1L2 (**Methods**). We were able to predict FI and FG levels in the DESIR data with the average R^2^ values of 0.30 and 0.17, respectively (Figure 4, **Supplementary Table 12**). The values were decreased as compared to those when trained and tested in the same data, i.e. NFBC1966 (R^2^ for FI=0.55, FG=0.41, **Supplementary Table 12**). RF and SVR-L2Linear-L1L2 produced a smaller change in the predictions (FI: RF=0.18, SVR-L2Linear-L1L2=0.22; FG: RF=0.21, SVR-L2Linear-L1L2=0.19), while BT was less stable in its performance (change in R^2^ 0.35 for FI, 0.32 for FG). Having a larger sample size in the training data (**Methods**) produced less variable R^2^ values than those resulting from the use of a smaller training dataset in all models (Figure 4A vs. 4B). While we still observed a drop in the R^2^ values for both FG and FI when tested in the external data, RF and SVR-L2Linear-L1L2 produced smaller decreases than BT, as before (**Supplementary Table 12**).

**Figure 4.**
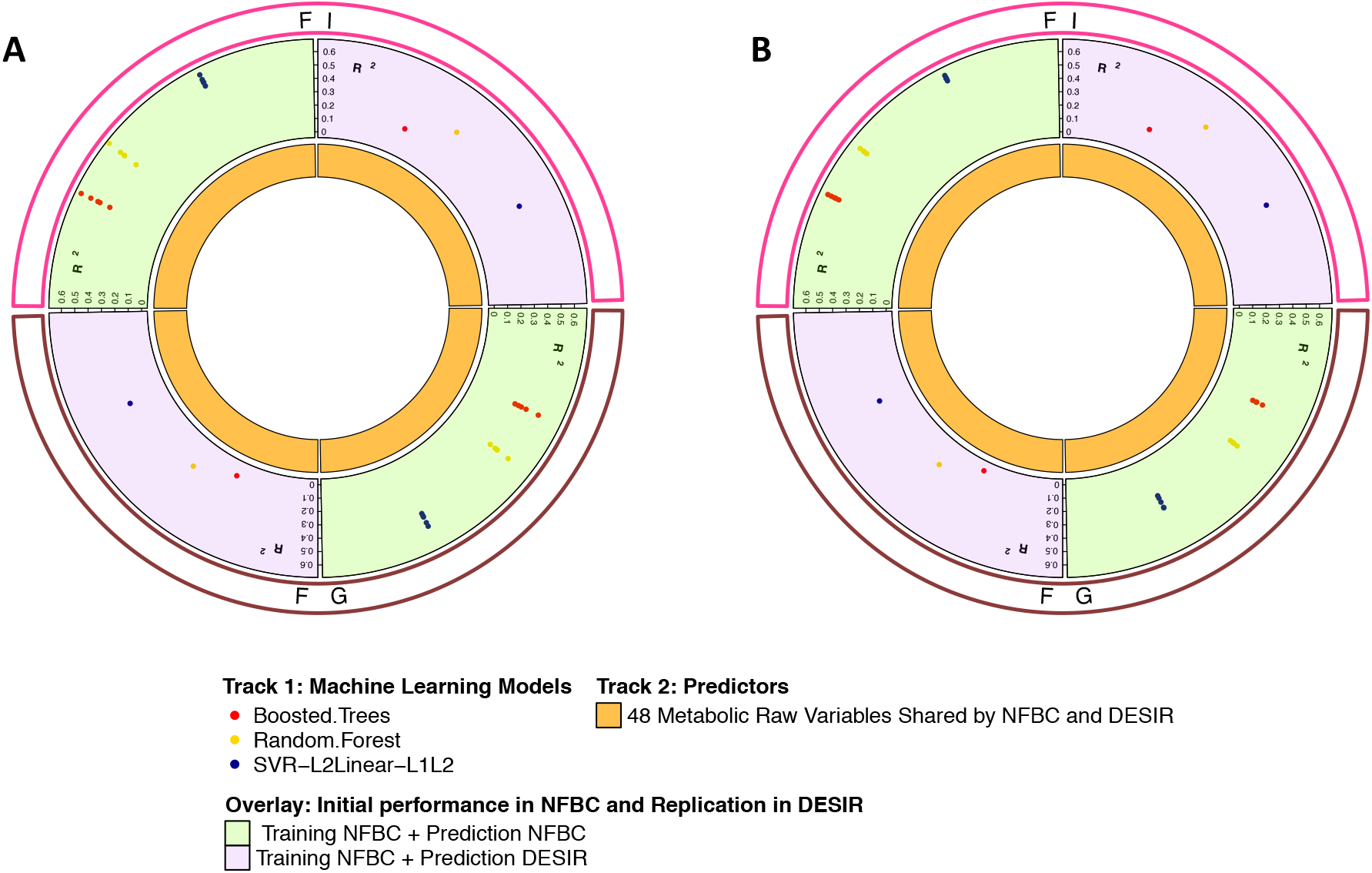
Performance as measured in R^2^ of the different machine learning models with 48 variables shared between NFBC1966 and DESIR. A. Using NFBC1966 data with individuals having methylation data (N=513) and all available DESIR individuals (N=769). B. Using NFBC1966 data with all available individuals (N=3,056) and all available DESIR individuals (N=769). Training of the algorithm was performed with a nested cross validation (5-folds outer, and 5-folds inner cross validation) and the R^2^ of 5 outer testing folds is displayed for each machine learning model. Metabolic predictors include epidemiological, biochemical and metabolomic data. SVR: Support Vector Regression with the kernel of linear function with L2 regularization and L1/L2 loss function (L2Linear-L1L2).

## Discussion

To our knowledge, this is the first multi-omics study implementing machine learning to predict continuous glycaemic traits over time. We dissected the predictive value of methylation and metabolic, including metabolomic, data from two time points for individual’s glycaemic health. We compared six machine learning approaches, while most previous studies have usually focussed on only one selected approach. We detected the best predictive ability of our models for FI and FG levels out of the five glycaemic traits tested, with raw or scored metabolic data as predictors and, BT, RF, or SVR-L2Linear-L1L2 as the algorithms. We identified metabolic variables that drove the prediction of the models. We showed that measures of adiposity are the most important contributors to glycaemic health. We also found that methylation probes accounted for 4 and 3 percentage points of the variance explained for FI (58%) and FG (45%), respectively. Finally, replication of the approach in an external European descent dataset (DESIR) using a subset of variables common to both cohorts, suggested that RF and SVR-L2Linear-L1L2 are more stable than BT in their performance.

Most of the published studies have targeted T2D onset prediction as a discrete value and rely on the categorization of individuals based on diagnosis thresholds for HbA1c, FG, 2hGluc and random glucose. The cut-off points for prediction analyses may vary across studies, and more generally, information loss is rather large when data is categorised^28,29^. Continuous phenotypes, on the contrary, have the potential to reflect the progressive onset of a disease without assuming a discontinuity in the underlying phenomenon. In addition, focussing on the prediction of continuous glycaemic phenotypes themselves allows removing them from the set of predictors for T2D and may reveal more modest effects of other variables^24^. Indeed, FG^23–27^ and 2hGluc^24,25^ have been shown to be good predictors of

T2D. Our study indicated that out of the five glycaemic traits we used, FG and FI were best predictable. This is expected as fasting values are tightly regulated. However, from the clinical practice point of view regarding the prediction of developing T2D, especially FI measurements have less relevance as compared to HbA1c measurements, for example. This suggests that future efforts should be directed towards improving the prediction of the other glycaemic indices than FG and FI.

Our study leverages machine learning ability to perform variable selection independently of a prefiltering. To date, RLS (a variant of SVR-L2Linear algorithms)^27^, J48-decision tree^25^, and logistic regression with regularization^23,26^ have highlighted the importance of specific metabolites consistent with our findings. Indeed, branched-chain amino acids (Leucine, Valine, Isoleucine)^25,26^, HDL, VLDL, glycerol, ApoA and Apo B, 3-hydroxybutyrate^26^, aromatic amino acids (phenylalanine, tyrosine)^23,26^ are established as important predictors by machine learning algorithms, as also shown by our study. Moreover, in this study, we report glycoprotein acetyls and acetoacetate as good predictors of glycaemic trait levels. These markers have previously been associated with T2D^30–32^; however, for the first time here, we show that they are not only associated, but are also predictors of glycaemic health.

In addition to specific metabolites, the machine learning algorithms assigned a high rank (first 25) to several established metabolic health-associated methylation probes in the prediction of FI and FG when collapsing metabolic predictors into scores but keeping methylation probes as such. The probes included for instance those within the genes *CPT1A* and *SREBF1*, where the first, *CPT1* (Carnitine palmitoyltransferase I) is involved in fatty acid metabolism (RefSeq, Jul 2008), and the latter, *SREBF1* (Sterol regulatory element-binding transcription factor 1) regulates genes required for glucose metabolism as well as fatty acid and lipid production, and its expression is regulated by insulin^33^. Previously, methylation at *CPT1A* and *SREBF1* has been associated with 2hIns^19^, BMI^20^, FG^18^ and T2D^18^. In our study, hypomethylation at *CPT1A* (cg17058475) at T2 predicted higher FI and FG levels, whereas hypermethylation at *SREBF1* (cg11024682) at T1 predicted higher FI levels. The predictive power of the methylation probes was modest, 0.04 out of 0.58 for FI and 0.03 out of 0.45 for FG. However, for both glycaemic traits the ranking of 15 methylation probes among the top 25 predictors when using scored metabolic data, sets the ground for larger studies of this kind, similarly to the work that has been achieved through large-scale GWAS. A recent study aggregating information over millions of genetic markers into a score showed that genetic risk scores for common diseases can identify people at risk equivalent to monogenic mutations^34^. On that account, the findings from our study encourage further exploration of methylation scores consisting of thousands or even millions of probes in glycaemic trait level prediction.

Measures of body adiposity and those of obesity are established risk factors for T2D^35^ and have a well-known impact on glycaemic trait variability^36^. In all six machine learning approaches and within all data combinations, we confirmed the high predictive value of BMI and WHR already 15 years beforehand. Indeed, when we calculated the variance explained from linear regression with the top 25 predictors, BMI and WHR accounted for the most part of it for both FI (0.40 out of 0.58) and FG (0.31 out of 0.45). When the outcomes were adjusted for the measures of adiposity, FG and FI levels at T1 gained more weight as predictors. These findings emphasize the importance of classical risk factors in T2D prediction but also show that the tracking is relevant already 15 years beforehand. Taken together, it is clear that classical risk factors will remain as valuable tools in the clinical practice for predicting future glycaemic health. However, more detailed biomarkers, for example certain metabolites as shown in the present study, genetic risk factors as shown recently^34^ and possibly methylation markers will open avenues for more precise prediction.

From the algorithm point of view, our analyses showed that the highest prediction performance was achieved with BT, RF and SVR-L2Linear-L1L2 algorithms. This is consistent with the literature as BT and RF have shown to perform well in the prediction studies of various types of data^37,38^. The SVR-L2Linear-L1L2 (LIBLINEAR library in R) algorithm performed significantly better than the SVR-L2LinearL1 (kernelab library in R). This difference can be explained by the use of the L2-loss over the L1-loss in the SVR-L2Linear-L1L2 in 78.5% of the cases (i.e. across all external cross-validations, all phenotypes and all data type combinations). We further investigated whether the better performance was due to the choice of the evaluation criterion, namely R^2^ which evaluates the performance in squared terms, similar to L2 loss, rather than in absolute terms, as do MAE and L1. Our analyses showed that SVR-L2Linear-L1 performed worse, regardless of the evaluation criterion used. These observations highlight the importance of loss function choice when using the SVR linear with L2-regularisation, independent of the model performance evaluation criterion. The SVR-Polynomial and SVR-RBF showed slightly lower predictions of the phenotypes, but there were no statistically significant differences when their performances were compared to those of the top three algorithms.

Our analysis has some limitations that warrant discussion. First, the relatively small sample size (513 subjects) is a drawback for taking full advantage of machine learning prediction with high-dimensional multi-omics data layers. However, our data is the largest to our knowledge featuring both longitudinal data and a comprehensive set of multi-omics data. UK Biobank has expressed its plans to acquire methylation data on its participants^39^. These data will be an important resource for future methylation studies. Nevertheless, the data will be cross-sectional and will not allow investigation of changes in methylation as the data used in the current study data does. Second, parameter tuning and drawing of a threshold regarding variable importance in our machine learning models are not trivial. Longer parameter tuning times might have resulted in more precise predictions and better performance of some or all of the algorithms. Variable importance in turn will depend on the number of variables resampled by the algorithms or the regularization parameters chosen. Overall, we restricted our analyses to six machine learning algorithms in total. It was out of the scope of this study to explore other potentially relevant algorithms, which will remain of future research interest. Third, regarding the samples and the study design, the use of whole blood only for methylation markers, and the relatively young age of the participants, 46 years old at the measurement time of the outcome variables is a limitation. The latter might alternatively represent a positive feature, since blood is the easiest tissue to obtain for any study, while the trend of deteriorating glycaemic health in younger adults is growing in all human populations. Finally, our replication effort also has some limitations. We were able to integrate only a small number of variables shared by both cohorts, and the time between the measurements differed between the Finnish and French studies, as did the age of the participants at recruitment. Despite these limitations the tested machine learning models showed promising consistency between the two cohorts, and we would expect better performance would the abovementioned limitations be addressed better.

## Conclusions

With the use of six different machine learning algorithms, we have identified clinically relevant sets of predictors of glycaemic traits from large multi-omics datasets and highlighted the potential of methylation markers and longitudinal changes in prediction. In the future, we expect that improvements in study sizes, methylation score computation, finer model tuning and replication in more similar external datasets will improve predictive ability of our models for glycaemic traits and will unveil novel prognostic omics biomarkers for T2D endophenotypes.

## Methods

### Study Populations

#### NFBC1966

Northern Finland Birth Cohort 1966 (NFBC1966) comprises participants from the two northernmost provinces of Finland with expected dates of birth falling in 1966 (N=12,058 births)^40^. From the medical examination at 31 (T1, N=6,007) and 46 years (T2, N=5,861) we included participants with demographic, medication, epidemiological, blood biochemical, metabolomic and epigenetic information available at both time points (N=626). Consent was obtained and the study was approved by the ethical committees of the University of Oulu and Imperial College London (Approval:18IC4421).

##### DESIR

The longitudinal DESIR study (Data from an Epidemiological Study on the Insulin Resistance syndrome) included 5,212 participants from the French general population. Clinical and biological evaluations were performed at inclusion and after three, six, and nine years, as previously described^41,42^. We used the data from 769 individuals aged from 30 to 65 years old at time of inclusion (T1) and nine years after inclusion (T2). Written informed consent was obtained from all participants. The study was approved by the Ethics Committee for the Protection of Subjects for Biomedical Research of Bicêtre Hospital, France.

### Epidemiological, Blood Biochemical and Metabolomic Data in the NFBC1966

Height, weight, waist and hip circumference, and systolic and diastolic blood pressure (SBP, DBP, each measured in triplicate) were measured according to standard study protocols at the clinical examinations at T1 and T2. We used the measured height and weight; however, if unavailable, data from postal questionnaire were used. Body mass index (BMI) was calculated from height and weight and waist-hip-ratio (WHR) from waist and hip measurements accordingly. The biochemical assays^43,44^, oral glucose^45^, and HbA1c measurements^46^ are detailed elsewhere. Metabolites were quantified by a high-throughput serum nuclear magnetic resonance (NMR) platform^47–50^. Imputation of epidemiological, biochemical and metabolomic variables was performed jointly with random forest (MisForest in R^51^) (**Supplementary Methods**). Post imputation, for individuals with diagnosed T2D at T2 (N=18), we corrected their potentially T2D medication-induced and thus artificially normal FG values to 7 mmol/l, HbA1c values to 48 mmol/l (6.5%) and 2hGluc values to 11.1 mmol/l. Detailed descriptions of all the exclusions and corrections are given in the **Supplementary Methods**. Finally, all the predictor variables were normalised using inverse-normal transformation. The epidemiological blood biochemical and metabolomic variables used in the predictions are listed in Table 1.

### Epigenomic Data in NFBC1966

DNA methylation was measured in whole blood from 807 randomly selected individuals after overnight fasting. At T2, DNA methylation was measured for 758 selected subjects that attended the clinical examination, completed the questionnaire and had DNA methylation data from the previous clinical examination available. IlluminaInfiniumHumanMethylation450 Beadchip and EPIC arrays were used at T1 and T2, respectively. Methylation data was quality controlled according to study protocol (**Supplementary Material**) and pre-processed on genome build CGCh37/hg19. Imputation of methylation data was performed with random forest (MisForest in R^51^) using the methylation residuals corrected for sex, and blood cell type (**Supplementary Material**). We limited our analysis to the methylation probes previously associated with seven phenotypes: 187 probes associated with BMI^20^, 21 with FG^18^, 11 with HbA1c^18^ and 68 with T2D^18^, one with 2hGluc^19^, eight with FI^19^, 21 with 2hIns^19^ (**Supplementary Table 2**).

### Epidemiological, Blood Biochemical and Metabolomic Data in DESIR

A total of 48 predictor variables overlapped between NFBC1966 and DESIR data. These included sex, measures of adiposity, biochemical data (triglycerides, total cholesterol, high and low-density lipoprotein cholesterol (HDL-C and LDL-C) at T1 and T2, insulin and glucose at T1), and metabolomic data (32 variables, 17 at T1 and 15 at T2). A full list of the included variables is given in **Supplementary Table 3**. The data were imputed with the package MisForest in R^51^ (missingness rate < 1%) (**Supplementary Material**). The values of FG were set to 7 mmol/l if the individual had diagnosed T2D. Finally, all the predictor variables were normalised using inverse-normal transformation.

### Individuals and Study Variables in the Machine Learning Models

In total, 513 individuals in the NFBC1966 were included in the machine learning analysis. HbA1c, 2hGluc, 2hIns, FG, FI levels were used as continuous outcomes to predict. A total of 1,001 variables from T1 and T2 were used as predictors in the NFBC1966. Metabolic predictors included: epidemiological data – sex, measures of adiposity (BMI and waist-to-hip ratio), SBP and DBP, biochemical data - ten blood measurements of triglycerides, total cholesterol, high and low-density lipoprotein cholesterol (HDL-C and LDL-C), metabolomic data - 454 metabolites (228 at T1 and 226 at T2) (**Supplementary Table 1**). The methylation dataset included 528 unique probes, including 264 at T1 and 264 at T2 (Supplementary Table 2).

For the replication analysis we chose to predict only FG and FI, for which the predictions worked the best in the NFBC1966 analyses. We included 48 variables common to both cohorts (**Supplementary Table 3**). The DESIR Cohort did not encompass methylation data. Therefore, before testing the prediction ability of our models in 769 individuals from DESIR, we could train the algorithms in the NFBC1966 either with the 513 individuals as previously, or with all 3,056 individuals who had metabolomics data available. This allowed us to test whether prediction in the external cohort was improved by increasing the sample size of the training data.

### Predictor Combinations and Prediction Frameworks

Metabolic (Mb) and Methylation (Mh) data were combined as their individual (or raw, denoted here as R) values or transformed into scores (S) (**Supplementary Material**). Methylation scores were formed according to the traits the probes have been previously associated with (**Supplementary Table 2**) and metabolic scores according to specific categories (**Supplementary Table 4**). We refer to Mb-R/Mh-R when the input data are all represented as individual values and to Mb-S/Mh-S when all the input data are combined in scores. Sex was kept separate and included in the model in addition to the scored variables. The following combinations were tested: Mb-R/ Mb-S/ Mh-R / Mh-S/ Mb-R + Mh-R/ Mb-R Mh-S/ Mb-S + Mh-R/ Mb-S + Mh-S (Figure 1). Methylation and Metabolic data were either adjusted for BMI and waist-hip-ratio at T1 and T2, or kept unadjusted.

### Machine Learning Approaches

Three machine learning methods were used for regression analysis: Boosted trees (BT), Random Forest (RF) and Support Vector Regression (SVR) (**Supplementary Material**). SVR was implemented with Linear Kernel with L2 regularization and with L1 and L1/L2 loss functions (SVR-L2Linear-L1, SVR-L2Linear-L1L2, respectively), with Polynomial Kernel (SVR-Polynomial) and with Radial Basis function Kernel (SVR-RBF) (Figure 1). For all the analyses, the packages kernelab, LIBLINEAR, RandomForest and xgboost in R^51^ were used with Caret as a wrapper.

### Optimization of the Machine Learning Algorithms

Nested cross validation was implemented. The data set was split into a training (80%) and testing set (20%) with a 5-fold cross validation. The performance of the machine learning models was estimated on the testing set, while parameter tuning was implemented on the training set by splitting it further into a 5-fold cross validation (nested). Random search method was used to find the model parameter combination (**Supplementary Table 13**) which minimized the error of the model. The Root Mean Square Error (RMSE) was used to assess model performance during training. Both Rsquared (R^2^) and RMSE were computed in the testing set to estimate performance. For additional checks we used the Mean Absolute Error (MAE) to assess model performance.

### Variable Importance in the Machine Learning Models

In BT, the information gain was used as a measure of importance. Gain is based on the decrease in entropy after a dataset is split on a feature j at a branch of the tree. RF variables were ranked with the Increase in Mean Square Error (MSE). It estimates the increase of prediction error when the values of the feature j are randomly permuted. For SVR, each feature is evaluated based on its independent association with the outcome. The slope of the regression is used to rank the features.

### Statistical Analysis for Comparing Model Performance

The performance of each model was computed as the average R^2^ over the 5 testing folds of the cross validation. In the Results section, we report the R^2^ pooled for the six machine learning algorithms. Comparison of the models was performed with a one-way ANOVA and post-hoc Tukey honest significance test (HSD) test. We use *P*-value<0.05 to denote statistical significance.

### Association Analysis to Obtain Effect Size, Direction and Variance Explained

We performed linear regression analyses between the best predicted outcomes (FG and FI) and each of the top 25 predictors suggested by the machine learning models to assess the effect sizes and directions of the associations. The analyses were conducted in R^51^. We report betas with their standard errors and related *P*-values. Additionally, we performed linear regression analyses for ln(FI)/FG by including all the top 25 predictors in the same model and then by removing metabolites/ methylation markers from the set of predictors to evaluate the variance explained and the contribution of metabolites/methylation markers in it. We report the adjusted R^2^ from these analyses.

### Replication analysis

For the replication analysis we used the three most consistently performing machine learning approaches: BT, RF, and SVR-L2Linear-L1L2. First, we estimated the performance of the algorithms in the NFBC1966 using a restricted set of 48 predictors. We selected the 48 input variables as overlapping with our validation cohort variables. This performance estimation was done by splitting the NFBC1966 dataset of 513 individuals into 80% for training and 20% for testing (nested cross validation for tuning as described above). Second, we “re-trained” the model with the maximum number of individuals in the NFBC1966, i.e. 513 individuals which represent 100% of the previous dataset with methylation and metabolomics data, or all 3,056 individuals with metabolomics data only. Then we tested for the ability of this model to predict accurately glycaemic traits when given an independent population (DESIR) with the same input variables. Practically, we used either 513 or 3,056 individuals from the NFBC1966 for training and used the resulting models to predict FI and FG in 769 individuals of the DESIR cohort. Performance was evaluated with R2 as described above.

## Supporting information

## Acknowledgements

IP is funded by the World Cancer Research Fund (WCRF UK) and World Cancer Research Fund International (2017/1641), the Wellcome Trust (WT205915), and the European Union’s Horizon 2020 research and innovation programme (DynaHEALTH, project number 633595). MAK works in a Unit that is supported by the University of Bristol and UK Medical Research Council (MC_UU_12013/1). The computational work was performed using the Imperial College Research Computing Service, DOI: 10.14469/hpc/2232.

We thank all the NFBC cohort members and researchers who participated in the 31 and 46 years study. We also wish to acknowledge the work of the NFBC project center. NFBC1966 31 years old study received financial support from University of Oulu Grant no. 65354, Oulu University Hospital Grant no. 2/97, 8/97, Ministry of Health and Social Affairs Grant no. 23/251/97, 160/97, 190/97, National Institute for Health and Welfare, Helsinki Grant no. 54121, Regional Institute of Occupational Health, Oulu, Finland Grant no. 50621, 54231. NFBC1966 46 years old study received financial support from University of Oulu Grant no. 24000692, Oulu University Hospital Grant no. 24301140, ERDF European Regional Development Fund Grant no. 539/2010 A31592.

The D.E.S.I.R. study has been funded by INSERM contracts with Caisse nationale de l’assurance maladie des travailleurs salariés (CNAMTS), Lilly, Novartis Pharma, and sanofi-aventis; INSERM (Réseaux en Santé Publique, Interactions entre les déterminants de la santé, Cohortes Santé TGIR 2008); the Association Diabète Risque Vasculaire; the Fédération Française de Cardiologie; La Fondation de France; Association de Langue Française pour l’Etude du Diabète et des Maladies Métaboliques (ALFEDIAM)/Société Francophone de Diabétologie (SFD); l’Office national interprofessionnel des vins (ONIVINS); Ardix Medical; Bayer Diagnostics; Becton Dickinson; Cardionics; Merck Santé; Novo Nordisk; Pierre Fabre; Roche; Topcon.

The D.E.S.I.R. Study Group. INSERM U1018: B. Balkau, P. Ducimetière, E. Eschwège; INSERM U367: F. Alhenc-Gelas; CHU D’Angers: Y Gallois, A. Girault; Centre de Recherche des Cordeliers, INSERM U1138, Bichat Hospital: F. Fumeron, M. Marre, R Roussel; CHU de Rennes: F. Bonnet; CNRS UMR8090, Lille: A. Fonnebond, S. Cauchi, P. Froguel; Centres d’Examens de Santé: Alençon, Angers, Blois, Caen, Chateauroux, Chartres, Cholet, Le Mans, Orléans, Tours; Institute de Recherche Médecine Générale: J. Cogneau; General practitioners of the region; Institute inter-Regional pour la Santé: C. Born, E. Caces, M. Cailleau, O Lantieri, J.G. Moreau, F. Rakotozafy, J. Tichet, S. Vol.

## Author contributions

IP conceived the study idea. IP, MK, LP, HD and ZB defined the phenotypes and quality control criteria. LP, HD, MDA, MW and MK performed the analyses. LY, BB, RR, SS, MAK, PF and MRJ provided the data. LP, MK and IP formed the central writing group. All authors participated in the critical revision of the manuscript and approved the final version.

## Competing Interests

The authors declare no competing interests.

## References

1. International Diabetes Federation – Home. Available at: https://www.idf.org/. (Accessed: 31st May 2018)

2. Buijsse, B., Simmons, R. K., Griffin, S. J. & Schulze, M. B. Risk assessment tools for identifying individuals at risk of developing type 2 diabetes. Epidemiol. Rev. 33, 46–62 (2011).

3. Abbasi, A. et al. Prediction models for risk of developing type 2 diabetes: Systematic literature search and independent external validation study. BMJ 345, 1–16 (2012).

4. Noble, D., Mathur, R., Dent, T., Meads, C. & Greenhalgh, T. Risk models and scores for type 2 diabetes: Systematic review. BMJ 343, 1243 (2011).

5. Herder, C., Kowall, B., Tabak, A. G. & Rathmann, W. The potential of novel biomarkers to improve risk prediction of type 2 diabetes. Diabetologia 57, 16–29 (2014).

6. Suhre, K. et al. Metabolic footprint of diabetes: a multiplatform metabolomics study in an epidemiological setting. PLoS One 5, e13953 (2010).

7. Fiehn, O. et al. Plasma metabolomic profiles reflective of glucose homeostasis in non-diabetic and type 2 diabetic obese African-American women. PLoS One 5, e15234 (2010).

8. Drogan, D. et al. Untargeted metabolic profiling identifies altered serum metabolites of type 2 diabetes mellitus in a prospective, nested case control study. Clin. Chem. 61, 487–497 (2015).

9. Scott, R. A. et al. An Expanded Genome-Wide Association Study of Type 2 Diabetes in Europeans. Diabetes 66, 2888–2902 (2017).

10. Mahajan, A. et al. Fine-mapping type 2 diabetes loci to single-variant resolution using high-density imputation and islet-specific epigenome maps. Nat. Genet. 50, 1505–1513 (2018).

11. Manning, A. K. et al. A genome-wide approach accounting for body mass index identifies genetic variants influencing fasting glycemic traits and insulin resistance. Nat. Genet. 44, 659–669 (2012).

12. Saxena, R. et al. Genetic variation in GIPR influences the glucose and insulin responses to an oral glucose challenge. Nat. Genet. 42, 142–148 (2010).

13. Soranzo, N. et al. Common variants at 10 genomic loci influence hemoglobin A1C levels via glycemic and nonglycemic pathways. Diabetes 59, 3229–3239 (2011).

14. Lowry, E. et al. Understanding the complexity of glycaemic health - Systematic bio-psychosocial modelling of fasting glucose in middle-age adults; a DynaHEALTH study. Int. J. Obes. In press, (2018).

15. Klose, R. J. & Bird, A. P. Genomic DNA methylation: The mark and its mediators. Trends Biochem. Sci. 31, 89–97 (2006).

16. Cortessis, V. K. et al. Environmental epigenetics: Prospects for studying epigenetic mediation of exposure-response relationships. Hum. Genet. 131, 1565–1589 (2012).

17. Chambers, J. C. et al. Epigenome-wide association of DNA methylation markers in peripheral blood from Indian Asians and Europeans with incident type 2 diabetes: a nested case-control study. Lancet. Diabetes Endocrinol. 3, 526–534 (2015).

18. Walaszczyk, E. et al. DNA methylation markers associated with type 2 diabetes, fasting glucose and HbA1c levels: a systematic review and replication in a case-control sample of the Lifelines study. Diabetologia 61, 354–368 (2018).

19. Kriebel, J. et al. Association between DNA Methylation in Whole Blood and Measures of Glucose Metabolism: KORA F4 Study. PLoS One 11, (2016).

20. Wahl, S. et al. Epigenome-wide association study of body mass index, and the adverse outcomes of adiposity. Nature 541, 81–86 (2017).

21. Hastie, T., Tibshirani, R. & Friedman, J. The Elements of Statistical Learning: Data Mining, Inference, and Prediction, Second Edition (Springer Series in Statistics). (2009).

22. Mühlenbruch, K., Jeppesen, C., Joost, H.-G., Boeing, H. & Schulze, M. B. The Value of Genetic Information for Diabetes Risk Prediction – Differences According to Sex, Age, Family History and Obesity. PLoS One 8, e64307 (2013).

23. Yengo, L. et al. Impact of statistical models on the prediction of type 2 diabetes using nontargeted metabolomics profiling. Mol. Metab. 5, 918–925 (2016).

24. Savolainen, O. et al. Biomarkers for predicting type 2 diabetes development-Can metabolomics improve on existing biomarkers? PLoS One 12, e0177738 (2017).

25. Allalou, A. et al. A Predictive Metabolic Signature for the Transition From Gestational Diabetes Mellitus to Type 2 Diabetes. Diabetes 65, 2529–2539 (2016).

26. Liu, J. et al. Metabolomics based markers predict type 2 diabetes in a 14-year follow-up study. Metabolomics 13, 104 (2017).

27. Peddinti, G. et al. Early metabolic markers identify potential targets for the prevention of type 2 diabetes. Diabetologia 60, 1740–1750 (2017).

28. van Walraven, C. & Hart, R. G. Leave ‘em Alone – Why Continuous Variables Should Be Analyzed as Such. Neuroepidemiology 30, 138–139 (2008).

29. Royston, P., Altman, D. G. & Sauerbrei, W. Dichotomizing continuous predictors in multiple regression: A bad idea. Stat. Med. 25, 127–141 (2006).

30. Bentley-Lewis, R. et al. Metabolomic profiling in the prediction of gestational diabetes mellitus. Diabetologia 58, 1329–1332 (2015).

31. Akbay, E. et al. The relationship between levels of alpha1-acid glycoprotein and metabolic parameters of diabetes mellitus. Diabetes. Nutr. Metab. 17, 331–335 (2004).

32. Mahendran, Y. et al. Association of ketone body levels with hyperglycemia and type 2 diabetes in 9,398 Finnish men. Diabetes 62, 3618–3626 (2013).

33. Ferre, P. & Foufelle, F. Hepatic steatosis: a role for de novo lipogenesis and the transcription factor SREBP-1c. Diabetes, Obes. Metab. 12, 83–92 (2010).

34. Khera, A. V et al. Genome-wide polygenic scores for common diseases identify individuals with risk equivalent to monogenic mutations. Nat. Genet. 50, 1219–1224 (2018).

35. Eckel, R. H. et al. Obesity and Type 2 Diabetes: What Can Be Unified and What Needs to Be Individualized? J. Clin. Endocrinol. Metab. 96, 1654–1663 (2011).

36. Scott, R. A. et al. Large-scale association analyses identify new loci influencing glycemic traits and provide insight into the underlying biological pathways. Nat. Genet. 44, 991–1005 (2012).

37. Caruana, R. & Niculescu-Mizil, A. An empirical comparison of supervised learning algorithms. Proc. 23rd Int. Conf. Mach. Learn. – ICML ‘06 161–168 (2006). doi:10.1145/1143844.1143865

38. Caruana, R., Karampatziakis, N. & Yessenalina, A. An Empirical Evaluation of Supervised Learning in High Dimensions. in International Conference on Machine Learning 96–103 (2008).

39. Grundberg, E. The opportunities of epigenomic research using UK Biobank data. Available at: http://www.ukbiobank.ac.uk/wp-content/uploads/2018/07/1405-Grundberg.pdf. (Accessed: 10th December 2018)

40. Northern Finland Cohorts. Available at: http://www.oulu.fi/nfbc/. (Accessed: 11th June 2018)

41. Vaxillaire, M. et al. Impact of common type 2 diabetes risk polymorphisms in the DESIR prospective study. Diabetes 57, 244–254 (2008).

42. Balkau, B., Eschwege, E., Tichet, J. & Marre, M. Proposed criteria for the diagnosis of diabetes: evidence from a French epidemiological study (D.E.S.I.R.). Diabetes Metab. 23, 428–434 (1997).

43. Taponen, S. et al. Hormonal Profile of Women with Self-Reported Symptoms of Oligomenorrhea and/or Hirsutism: Northern Finland Birth Cohort 1966 Study. J. Clin. Endocrinol. Metab. 88, 141–147 (2003).

44. Taponen, S. et al. Metabolic Cardiovascular Disease Risk Factors in Women with Self-Reported Symptoms of Oligomenorrhea and/or Hirsutism: Northern Finland Birth Cohort 1966 Study. J. Clin. Endocrinol. Metab. 89, 2114–2118 (2004).

45. Rautio, N. et al. Accumulated exposure to unemployment is related to impaired glucose metabolism in middle-aged men: A follow-up of the Northern Finland Birth Cohort 1966. Prim. Care Diabetes 11, 365–372 (2017).

46. Perkiömäki, N. et al. Association between Birth Characteristics and Cardiovascular Autonomic Function at Mid-Life. PLoS One 11, (2016).

47. Soininen, P. et al. High-throughput serum NMR metabonomics for cost-effective holistic studies on systemic metabolism. Analyst 134, 1781–1785 (2009).

48. Soininen, P., Kangas, A. J., Würtz, P., Suna, T. & Ala-Korpela, M. Quantitative serum nuclear magnetic resonance metabolomics in cardiovascular epidemiology and genetics. Circ. Cardiovasc. Genet. 8, 192–206 (2015).

49. Wurtz, P. et al. Quantitative Serum NMR Metabolomics in Large-Scale Epidemiology: A Primer on –Omic Technology. Am. J. Epidemiol. (2017).

50. Wang, Q., Holmes, M. V., Smith, G. D. & Ala-Korpela, M. Genetic support for a causal role of insulin resistance on circulating branched-chain amino acids and inflammation. Diabetes Care 40, 1779–1786 (2017).

51. R Core Team. R: A language and environment for statistical computing. (2014).

