## Supplementary material for "Machine Learning in Multi-Omics Data to Assess Longitudinal Predictors of Glycaemic Health"

### Supplementary Methods

**Phenotype imputation in the NFBC1966**

Prior to imputation, non-fasting (N=19) and pregnant (N=25) individuals at T1 were excluded. At T2, information on pregnancy was not available and none of the individuals were non-fasting. FI was log transformed to reduce skewness. Type 1 diabetes (T1D), T2D, gender, lipid lowering and diabetes medication were included as predictors in the imputation model. For individuals on blood pressure medication we added 15 mmHg and 10 mmHg to their recorded SBP and DBP values[1]. Data for men and women were imputed separately, since the use of oral contraceptives was added as a predictor for women in the imputation models at T1 (no information on contraceptive use at T2 was available).

Post-imputation, we excluded all T1D cases at both time points and additionally at T1, we excluded people with diagnosed T2D or on diabetes medication. For individuals with diagnosed T2D at T2, we corrected their FG values to 7 mmol/l, HbA1c values to 48 mmol/l (6.5%) and 2hGluc to 11.1 mmol/l. For lipid measures at both T1 and T2, we regressed out the effect of any lipid lowering medication and used the resulting residuals in the subsequent analysis. At both time points, we also calculated the averages of the three values for SBP and DBP and used these averages in the machine learning model. All measures of fasting/post-prandial glucose and insulin at T2 were removed from the set of predictor variables in the ML models as well as pyruvate at T2, which exhibited a high correlation with FG.

**Phenotype imputation in DESIR**

The measures of insulin at both time points were natural logarithm transformed. All measures of fasting glucose and insulin at T2 were removed from the set of predictor variables at both time points and pyruvate at T2 due to high correlation with the outcome variables. Data for men and women were imputed separately, similarly to the NFBC1966 analysis. All variables above were included in the imputation model. Imputation was performed with the package MisForest in R[2] (missingness rate < 1%). Differences in age at inclusion were corrected for by adjusting all the variables at T1 for age at T1 and using the resulting residuals in the machine learning analysis. The values of FG were set to 7 mmol/l if the individual had diagnosed T2D. No individuals had T1D in this study.

**Epigenomic data quality control in the NFBC1966**

For DNA methylation marker calling we used a detection *P*-value threshold of $<10^{-16}$ as suggested by Lehne *et al*[3]. A call rate filter of 95% was applied to all autosomal Illumina probes yielding 459,378 probes at T1 and 832,569 at T2. We removed duplicate samples (N=9/8 at T1/T2), gender mismatches (N=7/1 at T1/T2), samples with <95% call rate (N=67/40 at T1/T2) and with globally outlying DNA methylation values, i.e. 1st PC of the DNA methylation values outside mean +/- 4SD (N=1/1 at T1/T2). In total, 513 individuals were included in the machine learning analysis.
Intensity values were normalized with subset quantile normalization within array (SWAN in Minfi R[2]), and beta values were computed from methylated and unmethylated normalized probe intensities. Probes further than 4SD from the mean were removed. Batch and sex effect were corrected by including the principal components of the control probe intensities and gender as linear predictors in the regression analysis of the samples[4]. Blood cell composition was corrected for by using the Houseman estimates[5] of blood cell type in the regression.

**Calculation of Metabolic and Methylation Scores**

Unweighted risk scores were used. Metabolic risk scores at T1 and T2 grouped variables based on the following biochemical classes: lipoparticules, lipids, blood proteins, carbohydrates and insulin, keton-bodies, BRACA and other amino acids. Methylation scores at T1 and T2 were based on the established associations with seven phenotypes, including BMI, FG, HbA1c, T2D, 2hGluc, FI, 2hIns.

**Machine learning approaches**

*Decision trees*

We applied two decision tree -based methods, boosted trees (BT) and random forest (RF). The decision tree unit in both is a hierarchical framework. At each step the sample is split based on some threshold in one of the variables (feature). At each level, the feature examined must lead to the best possible prediction at the "leaves" level. Overfitting is avoided by a stopping criterion and recursive pruning of the tree. Boosting methods seek to construct iteratively predictors (e.g. trees, in BT) by focusing mis-predicted examples at the previous step[6]. RFs are an ensemble of decision trees. Each tree is grown on a bootstrapped sample of the data. The subset of features examined is generated randomly. The final prediction is based on the voting majority or averaging the predictions[6]. In RF, the growing of trees is done in parallel, as opposed to BT in which it is done iteratively. We used the xgboost implementation of BT[7–9] and RF[10]. All the analyses were performed with the packages xgboost and RandomForest in R[2] with Caret package as a wrapper.

*Support vector regression*

Support vector regression (SVR) is a regression technique based on an extension of support vector machine. SVR aims at fitting the data in a regression hyperplane by minimizing a margin. SVR allows for non-linearity in the data by mapping the data to a linear space via a Kernel function[6]. We applied SVR with Linear Kernel with L2 regularization and with L1 and L1/L2 loss functions (SVR-L2Linear-L1 and SVR-L2Linear-L1L2), with Polynomial Kernel (SVR-Polynomial) and with Radial Basis Function Kernel (SVR-RBF). The L2 regularization parameter in the linear SVRs allows handling of multicollinearity whereas the Polynomial and RBF kernels have the ability to handle non-linear relationships. All the analyses were performed in R[2] with Caret package as the wrapper. SVR-L2Linear-L1 analyses were performed with the package kernelab[11] in which the L1-loss refers to the epsilon-insensitive loss function of SVR; the sequential minimization optimization algorithm (SMO) is used for solving the SVR quadratic programming problem. SVR-L2Linear-L1L2 analyses were performed with the package LIBLINEAR[12]. In this case, the loss function (L1 or L2-loss) is an hyperparameter which choice is optimized during training. L2-loss is the squared version of the epsilon-insensitive loss function. LIBLINEAR uses a Newton-type method and a coordinate descent approach to solve the optimization problem[13]. The analyses for both SVR-Polynomial and SVR-RBF were performed with the kernelab[11] package.

*Parameter tuning*

Parameters were optimized as described in **Supplementary Table 12**. Random search was used for exploring the parameter space.

#### Supplementary References

1. Tobin, M. D., Sheehan, N. a., Scurrah, K. J. & Burton, P. R. Adjusting for treatment effects in studies of quantitative traits: antihypertensive therapy and systolic blood pressure. *Stat. Med.* **24**, 2911–2935 (2005).
2. R Core Team. R: A language and environment for statistical computing. (2014).
3. Lehne, B. *et al.* A coherent approach for analysis of the Illumina HumanMethylation450 BeadChip improves data quality and performance in epigenome-wide association studies. *Genome Biol.* **16**, 1–12 (2015).
4. Aslibekyan, S. *et al.* Association of Methylation Signals With Incident Coronary Heart Disease in an Epigenome-Wide Assessment of Circulating Tumor Necrosis Factor α. *JAMA Cardiol.* (2018). doi:10.1001/jamacardio.2018.0510
5. Houseman, E. A. *et al.* DNA methylation arrays as surrogate measures of cell mixture distribution. *BMC Bioinformatics* **13**, 86 (2012).
6. Hastie, T., Tibshirani, R. & Friedman, J. *The Elements of Statistical Learning: Data Mining,*

*Inference, and Prediction, Second Edition (Springer Series in Statistics)*. (2009).
7. Friedman, J., Hastie, T. & Tibshirani, R. Additive logistic regression: a statistical view of boosting. *Ann. Stat.* **28**, 337–407 (2000).
8. Friedman, J. Greedy function approximation: a gradient boosting machine. *Ann. Stat.* **29**, 1189–1232 (2001).
9. Friedman, J. H. Stochastic gradient boosting. *Comput. Stat. Data Anal.* **38**, 367–378 (2002).
10. Breiman, L. Random Forests. *Mach. Learn.* **45**, 5–32 (2001).
11. Karatzoglou, A., Smola, A., Hornik, K. & Zeileis, A. kernlab – An S4 Package for Kernel Methods in R. *J. Stat. Softw.* **11**, 1–20 (2004).
12. Fan, R.-E., Chang, K.-W., Hsieh, C.-J., Wang, X.-R. & Lin, C.-J. LIBLINEAR: A Library for Large Linear Classification. *J. Mach. Learn. Res.* **9**, 1871–1874 (2015).
13. Lin, C. Large-scale Linear Support Vector Regression. *J. Mach. Learn. Res.* **13**, 3323–3348 (2012).

#### Supplementary Figure and Table Legends

**Supplementary Figure 1. Performance as measured in $R^2$ of the different machine learning models after adjusting for measurements of adiposity (Waist-to-hip-ratio and Body mass index) at T1 and T2**. Training of the algorithm was performed with a nested cross validation (5-folds outer, and 5-folds inner cross validation) and the $R^2$ of 5 outer testing folds is displayed for each machine learning model. Metabolic predictors include epidemiological, biochemical and metabolomic data. SVR: Support Vector Regression with the kernels functions of linear with L2 regularization and L1 loss function (L2Linear-L1), linear with L2 regularization and L1/L2 loss function (L2Linear-L1L2), polynomial and radial basis function (RBF).

**Supplementary Table 1. Epidemiological, blood biochemical and metabolomic data at T1 and T2 in NFBC1966 and used for prediction of the levels of the five glycaemic traits.**

**Supplementary Table 2. Selected methylation probes, their availability and association analysis results between the probes and the phenotypes in the NFBC1966.**

**Supplementary Table 3. Variables common to both NFBC1966 and DESIR and used in the replication analysis.**

**Supplementary Table 4. Variables included in the different metabolic scores (Mb-S).**

**Supplementary Table 5. Results from the Tukey HSD test for comparing the outcome predictions according to input data. The predictions are averaged over all machine learning algorithms. Models are unadjusted for measurements of adiposity (Waist-to-hip-ratio and Body mass index) at T1 and T2.**

**Supplementary Table 6. Performance as measured in $R^2$ averaged over all machine learning models according to the input data.**

**Supplementary Table 7. Performance as measured in $R^2$ of all machine learning models according to the input data.**

**Supplementary Table 8. Results from the Tukey HSD test for comparing the different input data against each other in the prediction of the five glycaemic traits. The predictions are averaged over all machine learning algorithms. Models are unadjusted for measurements of adiposity (Waist-to-hip-ratio and Body mass index) at T1 and T2**

**Supplementary Table 9. Performance as measured in $R^2$ averaged over all machine learning models according to the input data after adjusting the outcomes for measurements of adiposity.**

**Supplementary Table 10. Results from the Tukey HSD test for comparing the performance of the different machine learning algortithms against each other in the prediction of the five glycaemic traits, according to different input data combinations. Models are unadjusted for measurements of adiposity (Waist-to-hip-ratio and Body mass index) at T1 and T2**

**Supplementary Table 11. Performance as measured in mean absolute error (MAE) of all machine learning models according to the input data.**

**Supplementary Table 12. Performance as measured in R2 of all the tested machine learning models when trained and tested on different sample sizes and on different cohorts.**

**Supplementary Table 13. Description of the parameters used for optimizing the machine learning models.**

#### Supplementary Figures

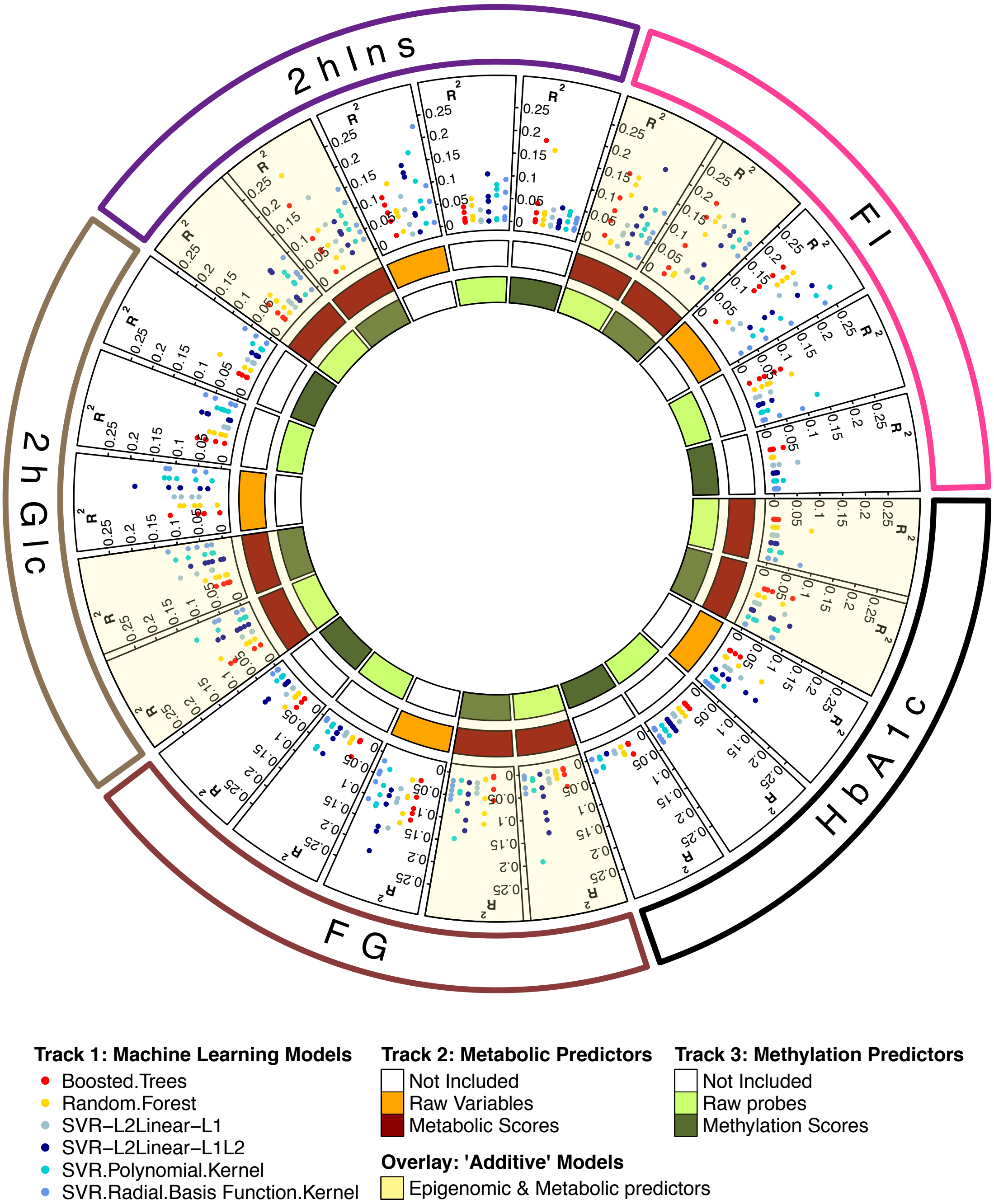

**Supplementary Figure 1. Performance as measured in $R^2$ of the different machine learning models after adjusting for measurements of adiposity (Waist-to-hip-ratio and Body mass index) at T1 and T2**. Training of the algorithm was performed with a nested cross validation (5-folds outer, and 5-folds inner cross validation) and the $R^2$ of 5 outer testing folds is displayed for each machine learning model. Metabolic predictors include epidemiological, biochemical and metabolomic data. SVR: Support Vector Regression (SVR) with the kernels of linear function with L2 regularization and L1 loss function (L2Linear-L1), linear with L2 regularization and L1/L2 loss function (L2Linear-L1L2), polynomial and radial basis function (RBF).

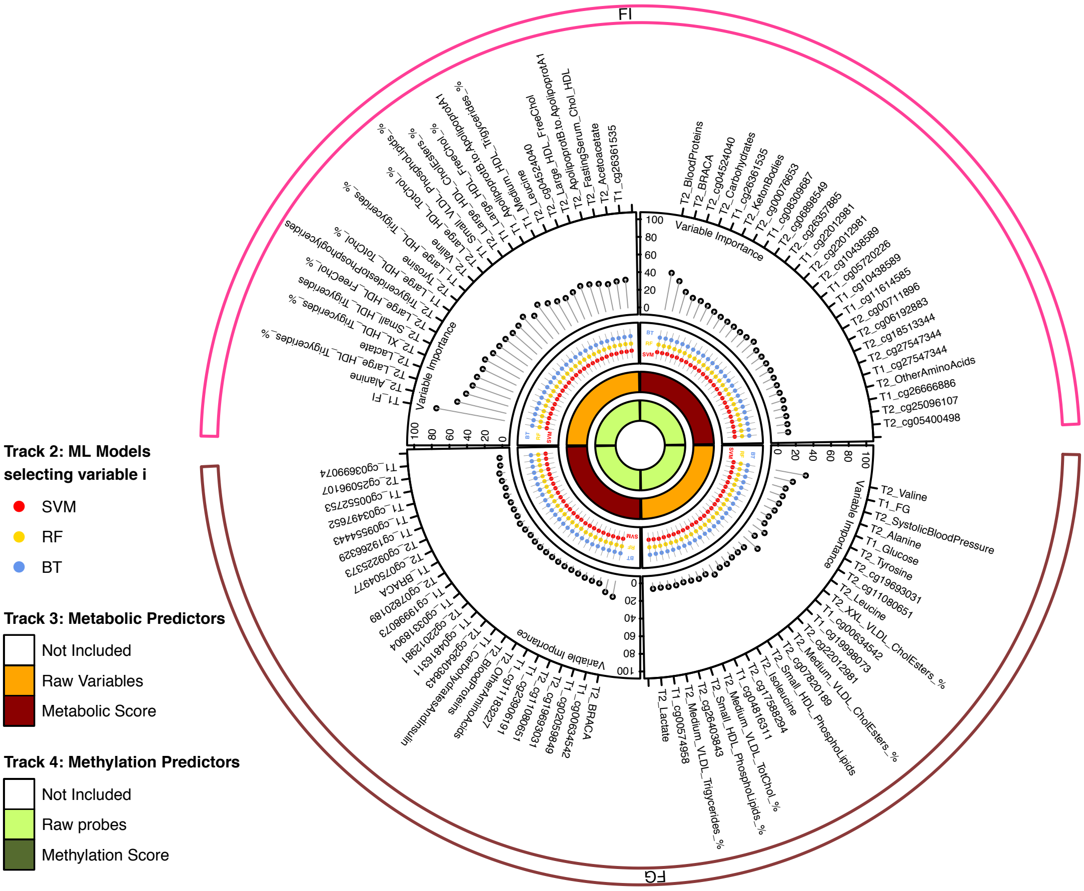

**Supplementary Figure 2. Variable Importance for fasting glucose and fasting insulin prediction from two different combinations of input data after adjustment for measurements of adiposity.** For each machine learning method, the normalized variable importance over five outer fold of cross validation was averaged into the "Variable-Model-Importance" (var.mod.Imp). Then for each of the six machine learning models, the variables were ranked based on the var.mod.Imp. The rank was averaged over the six models to obtain the mean variable rank. The latter was used to select top 25 variables for display. For these variables, we display the variable importance after (1) weighting the var.mod.Imp by the $R^2$ obtained for each of the individual ML algorithms (2) averaging variable importance across the six machine learning models. FI: Fasting Insulin; FG: Fasting Glucose; RF: random forest; BT: boosted trees, SVM: Support Vector Regression (SVR) with the kernels of linear function with L2 regularization and L1 loss function (L2Linear-L1), linear with L2 regularization and L1/L2 loss function (L2Linear-L1L2), polynomial and radial basis function (RBF). Metabolic predictors include epidemiological data, biochemical data and metabolomic data. T2: 46 years old, T1: 31 years old. BMI: Body Mass Index according to clinical examination, postal questionnaire if missing; WHR: Waist-to-hip ratio. Metabolite name descriptions are provided in **Supplementary Table 1**.

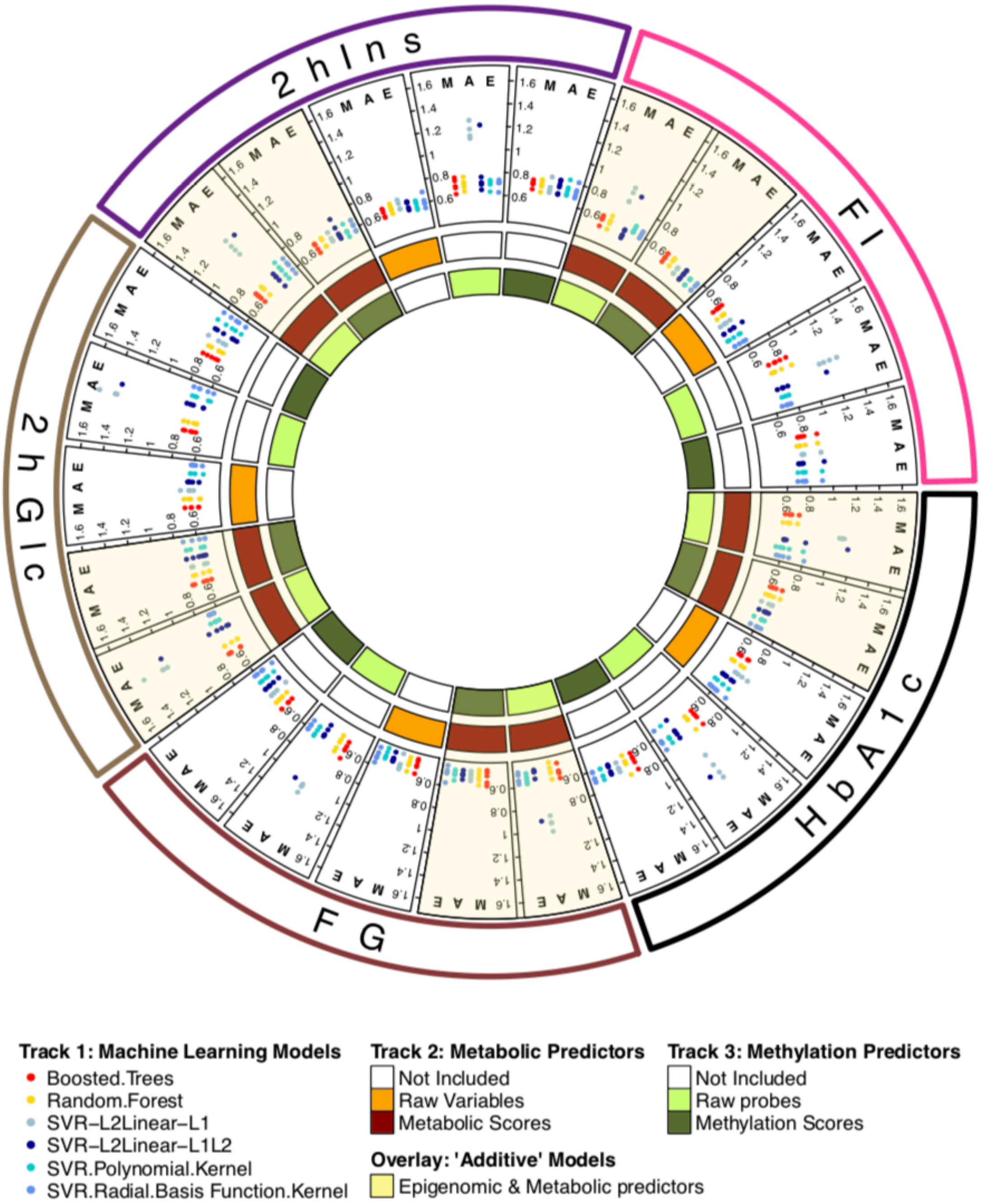

**Supplementary Figure 3. Performance as measured in MAE of the different machine learning models**. Training of the algorithm was performed with a nested cross validation (5-folds outer, and 5-folds inner cross validation) and the $R^2$ of 5 outer testing folds is displayed for each machine learning model. Metabolic predictors include epidemiological, biochemical and metabolomic data. SVR: Support Vector Regression (SVR) with the kernels of linear function with L2 regularization and L1 loss function (L2Linear-L1), linear with L2 regularization and L1/L2 loss function (L2Linear-L1L2), polynomial and radial basis function (RBF).
